# Investigating milk-derived extracellular vesicles as mediators of maternal stress and environmental intervention

**DOI:** 10.1101/2025.05.30.656911

**Authors:** Julia Martz, Baila Hammer, Tristen J. Langen, Benjamin Berkowitz, Benzion Berkowitz, Jasmyne A. Storm, Jueqin Lu, Deepali Lehri, Sanoji Wijenayake, Jordan Marrocco, Amanda C. Kentner

## Abstract

Parental communication signals are transmitted through nursing and critically shape neurodevelopmental trajectories. Mirroring some well characterized effects of gestational challenges in rodents, maternal immune activation (MIA) during the lactational period disrupts maternal physiology, decreases lipid content, and is associated with adverse neurobehavioral outcomes in offspring. This occurs without MIA significantly affecting maternal care. While gestational MIA models are responsive to environmental interventions, which beneficially alter maternal milk composition and associated offspring outcomes, the bioactive mediators in milk underlying resilience remain poorly understood. Milk-derived extracellular vesicles (MEVs) transport and deposit biologically active cargo, including microRNAs (miRNAs) that induce post-translational regulation of candidate mRNA in the nursing offspring’s tissues and cells. Using a rat model, we show that lactational MIA alters MEV-miRNA cargo and the expression of hippocampal miRNAs in offspring. Several miRNAs in MEVs were also found in the hippocampus of matching offspring. Remarkably, the miRNA changes in MEVs and the neonatal hippocampus were rescued when dams were raised in an enriched environment, suggesting environmental enrichment protected from the effects of MIA. This was supported by the behavioral phenotype. RNA-seq of adult offspring hippocampus showed long-term transcriptional changes associated with the gene targets of early-life regulated miRNAs. Our results position MEV-miRNA as dynamic programming signals by which maternal experience is communicated to offspring, encoding both stress-induced and protective cues that influence development. This suggests that breastfeeding interventions can regulate the genetic cargo of the milk, programming the life of developing infants.

## Introduction

Intergenerational transmission of biological cues shape offspring neurodevelopment and long-term behavioral outcomes. Beyond direct caregiving and nutritional transfer through nursing, recent research has identified extracellular vesicles (EVs) as a mechanism through which parental signals may be transmitted to offspring^1-7^. EVs are membrane-bound nanovesicles containing microRNAs (miRNAs), messenger RNAs (mRNAs), proteins, peptides, and lipids, capable of influencing gene expression and cellular function in target tissues^8-9^. While the mechanisms underlying parental contributions to offspring development via reproductive tract-derived EVs and sperm are well described^2,3^, maternal EV signaling during lactation remains less understood.

Breast milk is a biologically active medium; it supplies nutrition and other regulatory information to offspring^10^. Indeed, breastfeeding is associated with numerous benefits for infant development, including enhanced immune function, cognitive outcomes, and reduced risk for metabolic diseases^10-13^. Milk contains a substantial population of milk-derived extracellular vesicles (MEVs) that can survive the digestive tract, enter systemic circulation, and reportedly accumulate in peripheral tissues and the brain, including the hippocampus of nursing neonatal mice^5-6,14-19^. Complementary evidence from a blood brain barrier (BBB) cell-based model demonstrates that peripherally derived EVs transport their cargos across brain microvascular endothelial cells. This is most apparent under inflammatory conditions when barrier integrity is reduced^20^. Moreover, although the BBB is established in neonates, some brain associated barriers are more permeable to small molecules during this developmental stage^21-23^, a time when the brain shows increased bioavailability to peripherally administered EVs compared to adults^24^. These findings highlight that early life contexts characterized by reduced BBB permeability, whether developmental or inflammatory driven, may facilitate brain exposure to MEVs, providing a pathway for maternal signals to shape neurodevelopment.

EVs are increasingly recognized as important mediators of intercellular communication within the central nervous system (CNS). For example, EVs derived from both the periphery and CNS (e.g., neurons, astrocytes) can interact with and regulate microglia function^25-27^. A recent *in vitro* study demonstrated that MEVs are readily taken up by homeostatic and polarized human microglia, influencing the abundance and enzymatic activity of a critical epigenetic modulator, the DNA methyltransferase 1^27^. The study also illustrated a mechanistic association between DNMT1 regulation and MEV-miRNA 148a-5P. miRNAs are small noncoding RNAs that regulate post-transcriptional gene expression and affect neural development, primarily by acting as repressors of target messenger RNA^28,29^. Specific miRNA trafficked by MEVs are known to regulate synaptic development see ^15, 17,30^ and inflammatory responses, alongside other physiological functions^31^. Additionally, MEVs have beneficial effects in the gut through interactions with epithelial cells^14,32^ and may influence offspring health and development by directly influencing the microbiome via their miRNA cargos^6,18,32^. These findings suggest that MEVs have multiple points of contact with CNS target sites by which they can modify offspring neurodevelopment through postnatal transcriptional signaling.

Maternal infections during pregnancy are relatively common^34^ and have been linked to an increased risk of neurodevelopmental disorders in offspring^35^. To investigate the mechanisms underlying this association, animal models of maternal immune activation (MIA), where maternal immune stimulation during gestation induces neurodevelopmental impairments in offspring, are widely used and validated^36-37^. While these gestational MIA models are supported by epidemiological evidence and offer strong translational relevance, comparatively little is known about the prevalence or risks of infections during the lactational period^38^. Recently, animal models of MIA during lactation have been evaluated^39-41^, demonstrating that maternal immune challenges during the nursing period impact offspring neurodevelopmental outcomes similarly to the gestational models. Importantly, these behavioral effects occur despite minimal alterations in maternal care behaviors^39,40,42,43^. This suggests that other variables, like milk-borne factors, may mediate these outcomes. While stressors experienced across lactation can influence maternal physiology and milk composition^10,44^, comprehensive analyses of traditional milk factors such as nutritional content, immunoglobulins, inflammatory cytokines, and even the maternal immunogen used to stimulate lactational MIA have not fully explained the effects on offspring neurodevelopment^39^. Therefore, alternative mechanisms must be considered.

Given the capacity of MEVs to deliver regulatory molecules to recipient cells and impart biological effects^6,15,16,18^, and their demonstrated involvement in the CNS^27^, MEVs are a plausible candidate to mediate the effects of lactational MIA. Indeed, environmental experiences such as stress and illness modify the miRNA cargo of MEVs^7, 45-47^, supporting the hypothesis that MEVs are sensitive to maternal environmental conditions and may relay this information to nursing offspring. Based on these findings, we hypothesize that lactational MIA alters the cargo of MEVs, leading to changes in the transcriptomic profile of the developing brain. These molecular changes may drive long-term alterations in gene expression, contributing to behavioral abnormalities observed in adulthood. Furthermore, because environmental enrichment (EE) has been shown to mitigate the behavioral consequences of gestational MIA^48-50^ and to improve milk quality in healthy laboratory rats^51^, we propose that an EE intervention may stabilize MEV cargos, protecting against the effects of lactational MIA. Understanding these mechanisms will provide insight into how the maternal environment shapes offspring neurodevelopment via milk-derived signaling pathways and may suggest strategies to enhance resilience in at-risk populations.

## Methods and Experimental Overview

All animal protocols were approved by the MCPHS Institutional Animal Care and Use Committee and complied with AAALAC guidelines. The experimental timeline is illustrated in **Figure 1A**. Complete methodological details are provided in **Supplementary Methods, Supplementary Table 1**^37^, and DeRosa et al.^39,51^.

**Figure 1.**
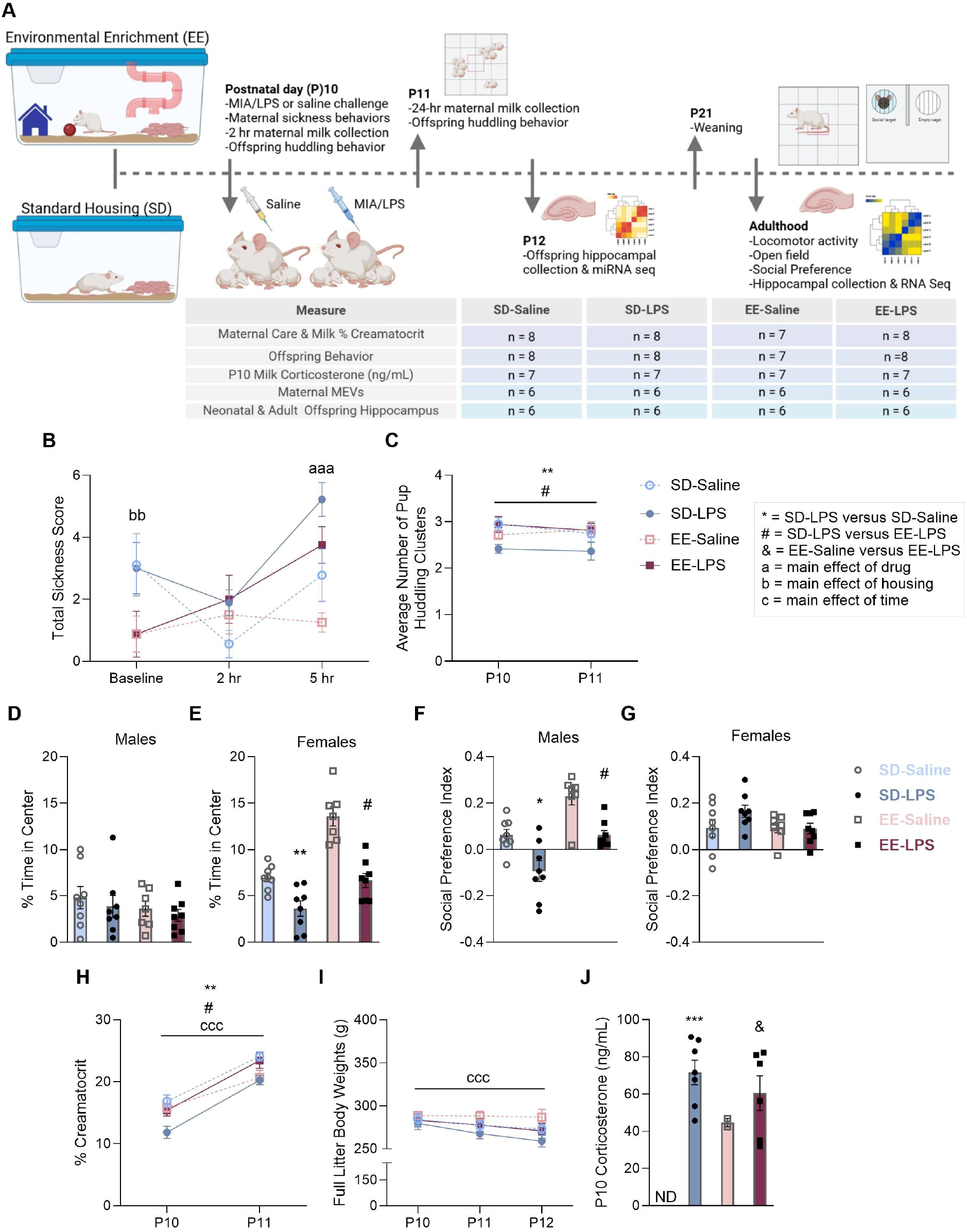
Environmental enrichment protects against lactational maternal immune activation (MIA) induced changes in offspring behavior. (**A**) Timeline of experimental procedures and group designations/sample sizes (n). (**B**) Maternal sickness behavior scores. **(C)** Neonatal huddling behavior (average number of pup clusters) across P10 and P11. Adult **(D)** male and (**E**) female percent (%) time in the center of the open field. Adult (**F**) male and (**G**) female social preference. (**H**) Percent (%) creamatocrit from P10-P11 and (**I**) the full litter body weights of nursing pups (g) across P10-P12. (**J**) Milk corticosterone (ng/mL) after MIA on P10; ND: non detectable values that fell below level of assay detection. Data are expressed as mean ± SEM; SD-Saline: n = 7-8; SD-LPS: n = 7-8; EE-Saline: n = 7; EE-LPS: n = 7-8. *p < 0.05, **p <0.01, ***p <0.001, SD-Saline versus SD-LPS; #p < 0.05, ##p <0.01, ###p <0.001, SD-LPS versus EE-LPS; ^&^p < 0.05, ^&&^p <0.01, ^&&&^p <0.001, EE-Saline versus EE-LPS; ^a^p < 0.05, ^aa^p < 0.01, ^aaa^p <0.001, main effect of drug; ^b^p < 0.05, ^bb^p < 0.01, ^bbb^p <0.001, main effect of housing; ^c^p < 0.05, ^cc^p < 0.01, ^ccc^p <0.001, main effect of time; LPS: lipopolysaccharide; P: postnatal day. *Created in BioRender. Kentner, A. (2025) https://BioRender.com/jrb10y6*.

### Animals and Housing

Sprague Dawley rats (Charles River, Wilmington) were bred and housed in either environmentally enriched (EE) or standard (SD) laboratory conditions as previously described^49,51^. On postnatal day (P)10, lactating dams were administered either 100 μg/kg lipopolysaccharide (LPS; *Escherichia coli* O26:B6) or pyrogen-free saline via intraperitoneal injection (n = 7-8 litters/group). Dams were separated from their litters to allow milk to accumulate for 2 hours post-injection while litters were kept warm on a heating pad^39,51^. Based on previous work, this period of separation does not induce a strong corticosterone response in neonates^52^.

### Maternal Assessment

Sickness behavior was evaluated at baseline, 30-, 120-, and 300-minutes post-injection using a composite score of ptosis, piloerection, and lethargy (0 = none, 1 = mild, 2 = severe), by blinded investigators, as adapted from prior studies^48,53,54^. Maternal care was observed on P9–P11 during AM and PM sessions. Behavioral metrics included pup licking/grooming frequency and time spent on the nest across six 1-minute intervals^39,51^.

### Milk Collection and Analysis

Two hours after injection, dams were anesthetized with isoflurane and administered oxytocin (0.2 mL; 20 USP/mL, i.p.) to induce milk letdown^39,51^; based on their pharmacokinetic profiles, these drugs are not absorbed by offspring^55-57^. Approximately 2.0 mL of milk was manually expressed and collected over 50–60 minutes. Dams were returned to their litters once recovered from anesthesia. Milk was collected again 24 hours later on P11. Percent creamatocrit was determined following procedures previously outlined^39,51, 58^, and 100 µL of milk was stored at –80L°C for corticosterone assays. Samples were thawed and rotated overnight, then analyzed using a small sample ELISA protocol (#ADI-900–097, Enzo Life Sciences; 1:40 dilution)^39,51^.

### Milk-Derived Extracellular Vesicles (MEVs)

Following centrifugation to remove the cream layer, MEVs were isolated from the supernatant using differential ultracentrifugation and filtration (DUC), as previously outlined^2, 3, 27,59^ **(Figure 2A)** and compliant with MISEV 2023 guidelines^60^. Casein was precipitated^61^, and whey fractions were ultracentrifuged to pellet MEVs, which were washed and resuspended in PBS. MEVs were characterized per MISEV 2023 guidelines using nanoparticle tracking analysis (NTA), western blotting, and transmission electron microscopy (TEM). NTA determined MEV size and concentration (**Figure 2B**). Western immunoblotting confirmed the presence of markers CD-9, Alix, Flotillin-1, and the absence of Calnexin (**Figure 2C**); these protein markers were assessed qualitatively to verify MEV identity^27,59,60^. TEM verified MEV morphology (**Figure 2D**).

**Figure 2.**
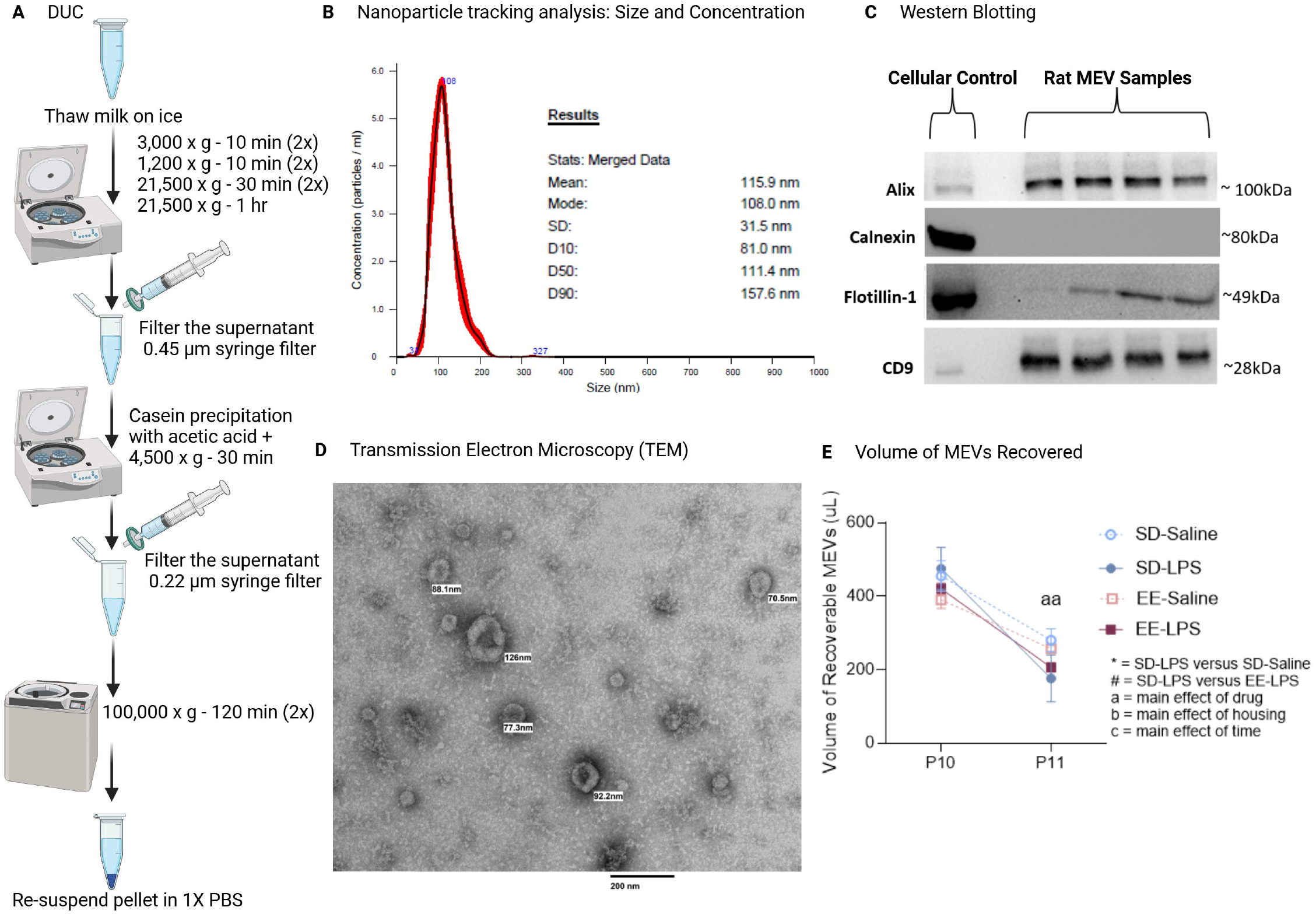
Characterization of milk-derived extracellular vesicles (MEVs) isolated from rat milk following maternal immune activation (MIA). (**A**) The differential ultracentrifugation (DUC) and serial filtration protocol followed for MEV isolation. (**B**) Size (nm) and concentration (particles/mL) of MEVs as determined by nanoparticle tracking analysis (NTA) with a mean size of 115.9 ± 31.5 nm. (**C**) The presence or absence of positive (Alix, Flotillin-1, CD9; cellular control: human MEVs) and negative (Calnexin; cellular control: Hep G2 cells) protein markers for MEVs as determined by western immunoblotting. **(D)** Transmission electron microscopy (TEM) of a representative MEV pellet. Scale bar is 200 nm with some individual MEVs identified in sizes ranging from 70.5 to 126 nm. (**E**) Total yield of MEVs recovered. Data are expressed as mean ± SEM, n = 6. ^aa^p < 0.01, main effect of drug; LPS: lipopolysaccharide; P: postnatal day; SD: standard housing. *Created in BioRender. Kentner, A. (2025) https://BioRender.com/x9rk66v*.

### Offspring Assessments

The offspring of saline and MIA dams were evaluated on neonatal huddling behavior (a proxy for neonatal thermoregulation; n = 7-8 litters/group), two hours after reuniting with their dams on P10 and P11^39,62^. Litters were weaned into same-sex pairs on P21, maintaining their original housing assignments. Starting on P70, one male and one female offspring from each litter were assessed on the open field and social preference tests (n = 7-8 litters/group/sex)^39,49,63^. One male and one female from each litter was euthanized with isoflurane, and whole (at P12) or ventral^64-66^ (at P74) hippocampus dissected, frozen on dry ice and stored at −80°C.

### RNA Extraction and Sequencing

Total RNA was extracted from hippocampal tissues using the RNeasy Lipid Tissue Mini Kit (74804, Qiagen) and from MEV samples using the Exosomal RNA Isolation Kit (58000, Norogen Biotek), according to the manufacturer’s instructions. Samples were quantified using Qubit 2.0 Fluorometer (ThermoFisher Scientific) and RNA integrity was checked with 4200 TapeStation (Agilent Technologies).

#### MEV small RNA-seq (n = 6 litters/group)

Libraries were prepared using NEB Small RNA Library Prep Kit (New England Biolabs), validated on TapeStation, quantified using Qubit 2.0 Fluorometer, and sequenced on Illumina NovaSeq (2×150bp PE), and de-multiplexed using Illumina’s bcl2fastq 2.20 software.

#### Neonatal hippocampus small RNA-seq (n = 6 litters/group)

Libraries were prepared using Illumina TruSeq Small RNA Library Prep Kit, cDNA constructs purified via BluePippin, validated on TapeStation, quantified using Qubit 2.0 Fluorometer, and sequenced on Illumina NextSeq 2000 (1×50bp).

#### Adult hippocampus RNA-seq (n = 6 litters/group)

Strand-specific libraries were generated using NEBNext Ultra II Directional RNA Library Prep Kit (Illumina), validated on TapeStation, quantified using Qubit 2.0 Fluorometer, and sequenced on Illumina NovaSeq (2×150bp PE), and de-multiplexed using Illumina’s bcl2fastq 2.20 software.

### Sequencing Analysis

Differentially expressed miRNAs and genes were identified based on a p<0.01, (FC)>2.0. Heatmaps were generated using Multiple Experiment Viewer. Venn diagrams were generated using DeepVenn (https://www.deepvenn.com). Gene ontology was determined using the DAVID functional annotation cluster tool (https://david.ncifcrf.gov/), while miRNA Enrichment Analysis and Annotation Tool (miEAA) was used to functionally annotate the different sets of miRNAs. Bubble plots and pie charts were generated in GraphPad Prism 10.4.2.

### Statistical Analyses

Data were analyzed using Prism (GraphPad) or SPSS (IBM). ANOVAs (Housing x MIA x Time or Housing x MIA) were used to evaluate milk % creamatocrit and behavioral endpoints, unless there were violations to the assumption of normality (Shapiro-Wilk test) in which case Kruskal-Wallis tests were employed (expressed as *X*^*2*^). To assess differences in the proportion of samples with detectable P10 milk corticosterone concentrations, the Fisher-Freeman-Halton exact test^67^ was used to appropriately account for the undetectable hormone levels in the SD-Saline group. Partial eta-squared (ηp^2^) is reported as an index of effect size for the ANOVAs^68^. Both male and female animals were included, and separate analyses were run for each sex^39,51,68,69^. All data are expressed as mean ± SEM.

## Results

### EE protected against lactational MIA-induced changes in offspring behavior

Maternal immune challenge during lactation significantly increased sickness behaviors in dams, regardless of housing assignment, validating the model. A MIA by time interaction identified an increased severity of sickness behavior in LPS treated dams 5 hours post immune challenge, compared to saline dams (**Figure 1B**). A housing by time interaction indicated that SD dams appeared less healthy than EE dams at baseline, consistent with prior findings^51^. SD dams, confined to smaller cages, nurse more due to a limited ability to escape from pups. SD housing is associated with displays of “pressing” behavior where dams press their ventral side against the cage, seemingly to hide their teats and take a break from the energy-intensive nursing task^51^.

Huddling behavior was significantly reduced in SD-LPS neonatal offspring compared to neonates from SD-Saline dams (MIA by housing interaction; average number of clusters **Figure 1C**). Diminished neonatal huddling behavior was buffered by EE (SD-LPS versus EE-LPS).

In the open field test, MIA by housing interactions were observed for both adult male and female offspring in terms of distance traveled (**Supplementary Figure 1A, B**). However, follow up tests were not significant in males; female EE-LPS rats traveled less than SD-LPS females. While males were not affected, SD-LPS female offspring spent a reduced percentage of time in the center of an open field compared to SD-Saline females (MIA by housing interaction: **Figure 1D, E**). This MIA induced avoidance behavior was prevented by EE housing. In contrast, male SD-LPS adult offspring had a decreased social preference index compared to SD-Saline males (MIA by housing interaction: **Figure 1F, G**). Complex EE housing also protected against the MIA associated alteration in male social behavior.

Maternal care differences do not appear to account for these effects since there were no significant group differences in terms of total time on nest or the number of pup-directed licking or grooming bouts (**Supplementary Figure 1C, D**). While previous work has shown housing differences in the maternal care of untreated rats^51^, stress induced by the LPS/Saline injections may have normalized the amount of maternal care displayed across SD and EE groups in the present study. **Statistics reported in Supplemental Table 3**.

### Housing condition regulates lactational MIA-induced changes in MEV-miRNA cargos

Building on our previous observations that MIA reduces milk quality^39^ while EE improves it^51^, we investigated whether EE housing could mitigate MIA-induced changes to milk composition. Percent creamatocrit was decreased in the milk of SD-LPS dams compared to SD-Saline, which was mitigated by EE housing (MIA x Housing interaction; main effect of time; **Figure 1H**). However, the body weights of nursing pups were not affected by either MIA or housing across P10-P12 (p>0.05; **Figure 1I**) nor at weaning (p>0.05; **Supplemental Figure 1E**). Milk corticosterone, known to influence offspring development and temperament^71,72^, differed significantly across the experimental groups (p = 0.001). Specifically, milk corticosterone was detectable following MIA on P10 (SD-Saline versus SD-LPS: p = 0.001; EE-Saline versus EE-LPS: p = 0.015) which was not reduced by EE (SD-LPS versus EE-LPS: p > 0.05; **Figure 1J**). Therefore, it is unlikely that attenuation of corticosterone underlies the protective effects of EE on behavior. **The full statistical results are reported in Supplemental Table 3**.

Although the total yield of MEVs recovered during the isolation process was reduced by MIA on P11 (MIA x Time interaction; **Figure 2E**) this was also not protected by EE in this model. Given the characteristics of MEVs to carry bioactive cargos such as miRNAs^15^, we turned our interests towards the effects of MIA and EE on MEV cargo composition. The cargo of MEVs isolated from SD-LPS dams showed 66 differentially expressed miRNAs compared to MEVs from SD-Saline dams. Specifically, 22 miRNAs were differentially expressed at P10, while 46 miRNAs were differentially expressed at P11, indicating that it takes about 24 hours for peak MEV-miRNA changes to appear (**Figure 3A,C**). Curiously, in the cargo of MEVs isolated from EE-LPS dams only 29 miRNAs, of which 16 at P10 and 14 at P11, were differentially expressed compared to MEVs from EE-Saline dams (**Figure 3B,D**). This means that, when dams were raised in EE, the number of LPS-induced miRNAs did not vary the day after the LPS challenge (P11) as opposed to what was observed in dams raised in SD (**Supplemental Figure 2**). In addition, we found that novel miRNA-629 was downregulated both at P10 and P11 in SD-LPS MEVs compared to SD-Saline MEVs, while the regulation of novel miRNA-375 diverged between P10 and P11. Finally, novel miRNA-1347 was upregulated in the MEVs of LPS-treated dams compared to Saline-treated dams raised in EE, regardless of timepoint (**Figure 3C,D**). Together, these data highlight the protective effects of EE on MEV-miRNAs both acutely and chronically, given the sustained ability of enrichment to regulate this cargo.

**Figure 3.**
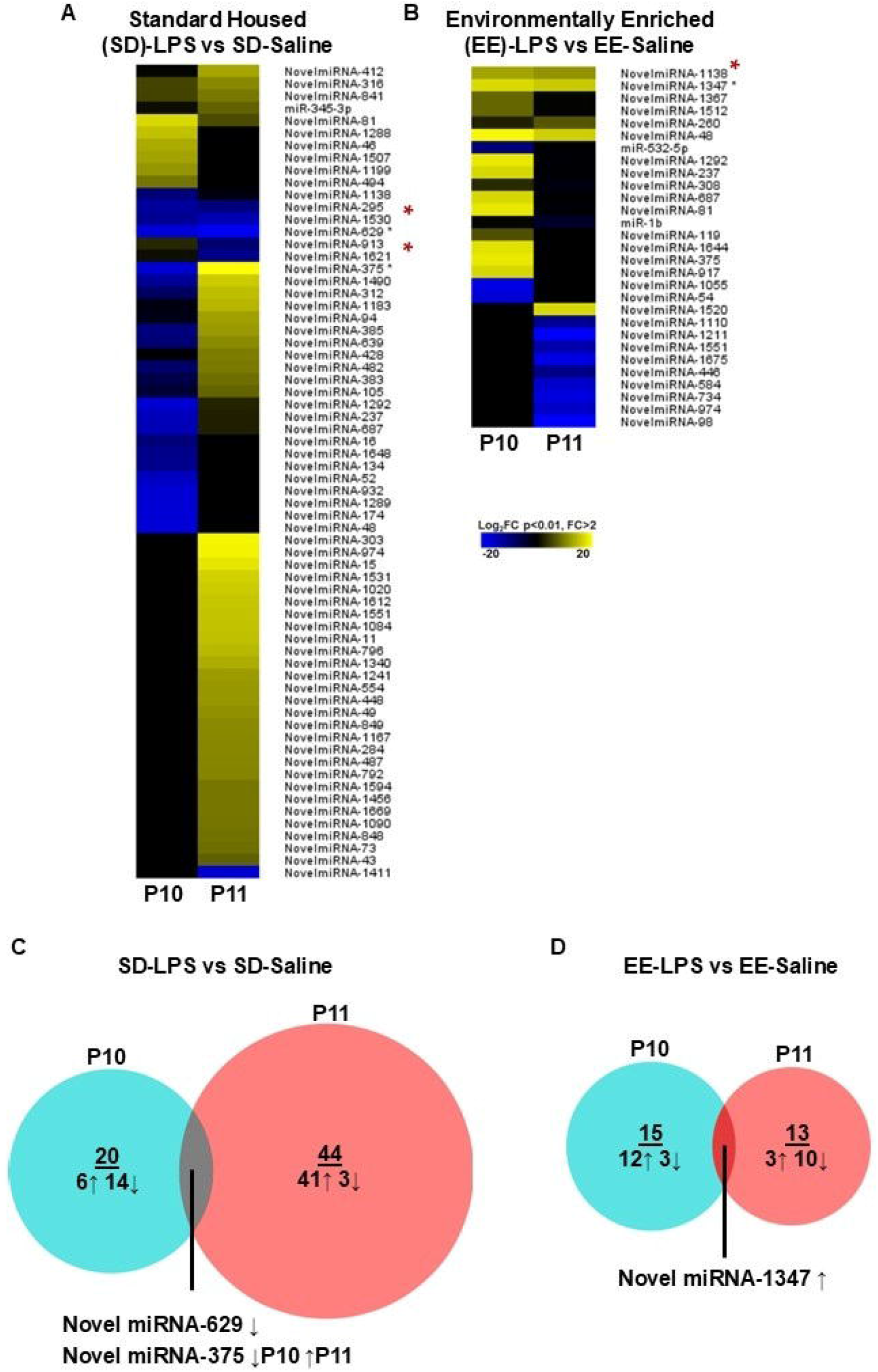
Lactational maternal immune activation (MIA) alters milk-derived extracellular vesicles (MEV) miRNA cargos, which are buffered by environmental enrichment (EE) housing. miRNA clustered in the heatmaps display the Log_2_FC p<0.01, FC>2 in (**A**) SD-LPS versus SD-Saline and (**B**) EE-LPS versus EE-Saline MEVs at P10 and P11. Venn diagrams show the number of differentially expressed miRNA in P10 (blue) and P11 (red), and their overlapping miRNAs and direction of change (↑ upregulated, ↓ downregulated) for (**C**) SD-LPS versus SD-Saline and (**D**) EE-LPS versus EE-Saline. LPS: lipopolysaccharide; P: postnatal day; SD: standard housing; n = 6. *If significant at P10 and P11.

### The composition of miRNAs in the nursing offspring hippocampus correlates with MEV-miRNA cargo as a function of housing

We then sought to explore the expression of miRNAs in the hippocampus of the nursing offspring (P12) of dams whose MEV-miRNA cargo was significantly affected by MIA as a function of EE. LPS caused differential expression of 50 miRNAs (45 miRNAs, adj p-value <0.05) in SD males and 4 miRNAs (miR-1247-5p, adj p-value<0.05) in SD females (**Figure 4A**). In contrast, as also observed in MEVs from EE dams, EE dramatically reduced the number of differentially expressed LPS-related miRNAs in the hippocampus compared to miRNAs induced by LPS in SD rats. Namely, LPS regulated 2 miRNAs in EE males and 5 miRNAs in EE females (2 miRNAs, adj p-value <0.05) compared to their respective Saline-treated EE controls (**Figure 4B**).

**Figure 4.**
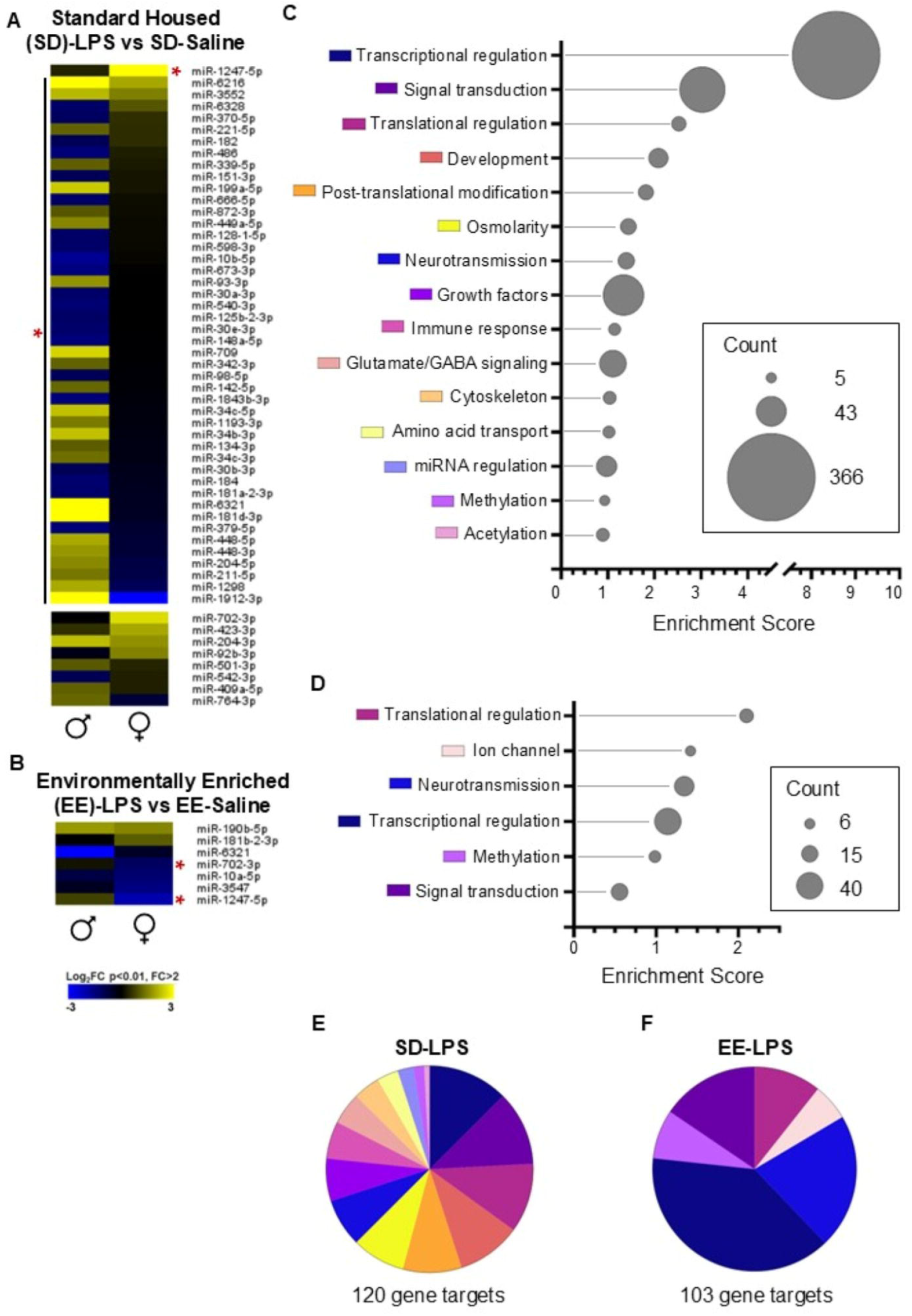
Lactational maternal immune activation (MIA) alters miRNA expression in the hippocampus of nursing offspring, which is buffered by environmental enrichment (EE) housing. miRNA clustered in the heatmaps display the Log_2_FC p<0.01, FC>2 in (**A**) SD-LPS versus SD-Saline and (**B**) EE-LPS versus EE-Saline neonatal hippocampus on P12 for male and female offspring. (**C,D**) The bubble plots show the enrichment score of gene pathways generated using DAVID by clustering genes predicted as miRNA-mRNA targets (mirdb.org, target score >80). Gene counts per cluster are represented as bubble size. Gene pathways highlighted are top unique clusters from (**C**) Figure A, and (**D**) Figure B. (**E,F**) Pie charts to visualize gene count distribution of pathways highlighted in (**E**) Figure C and (**F**) Figure D. LPS: lipopolysaccharide; P: postnatal day; SD: standard housing; n = 6; *if FDR<0.05. Edited in BioRender. Kentner, A. (2025) https://BioRender.com/w5acrdr

We were interested in understanding the genomic function of differentially expressed miRNAs regulated by LPS as a function of housing across groups. To predict the miRNA-mRNA targets, we used miRDB (threshold target score>80) combined with DAVID to cluster predicted genes according to their gene ontology. In selected pathways of interest, we found that miRNA-targeted genes in LPS rats (120 in SD rats and 103 in EE rats) were involved in several cellular functions including transcriptional and translational regulation, signal transduction, neurotransmission, as well as epigenetic regulation, regardless of their housing condition. Notably, glutamatergic/GABAergic signaling, immune response, growth factors, and development-related pathways, were targeted in LPS rats raised in SD, but not in EE (**Figure 4C-F**).

Building on the evidence that LPS caused behavioral alterations in adulthood that depended on early housing condition, we investigated the genomic profile in the ventral hippocampus of the adult offspring. The downregulation of *Thbs1*, a gene involved in autism spectrum disorder risk^73,74^, in the hippocampus of male rats whose mothers were raised in SD and challenged with LPS, was consistent with the upregulation of the miRNA regulator, miR-709, in the hippocampus at P12. The same was true for the upregulation of the gene *Naa11*, also involved in autism susceptibility^75^, whose regulator, miR-128-1-5p, was downregulated at P12 in the hippocampus of male rats whose mothers were raised in SD and challenged with LPS. We found that miRNA targets at P12 predicted the regulation of several genes that belonged to the same gene superfamily of the ones that were differentially expressed in the adult ventral hippocampus (**Supplemental Table 2**).

To assess whether miRNAs differentially expressed in MEVs were also affected in the hippocampus of the nursing offspring at P12, we compared the miRNA counts in both miRNA-seq datasets. While the overall pattern of common expression between MEV and hippocampus did not depend on treatment, housing, or sex, there were differences in the genomic function of miRNA sets across groups. Using miRNA Enrichment Analysis and Annotation Tool, we found that, out of the top five pathways, integrin and dopamine signaling, as well as adrenalin and noradrenalin biosynthesis, were regulated across all experimental groups (**Figure 5A-H**). However, the regulation of the glutamate signaling pathway was exclusive to LPS-treated groups raised in EE (**Figure 5G,H**). Additionally, the drug metabolism P450 pathway was regulated in all groups except EE-LPS. Again, this suggests that EE has unique effects on LPS-induced transcriptional regulation.

**Figure 5.**
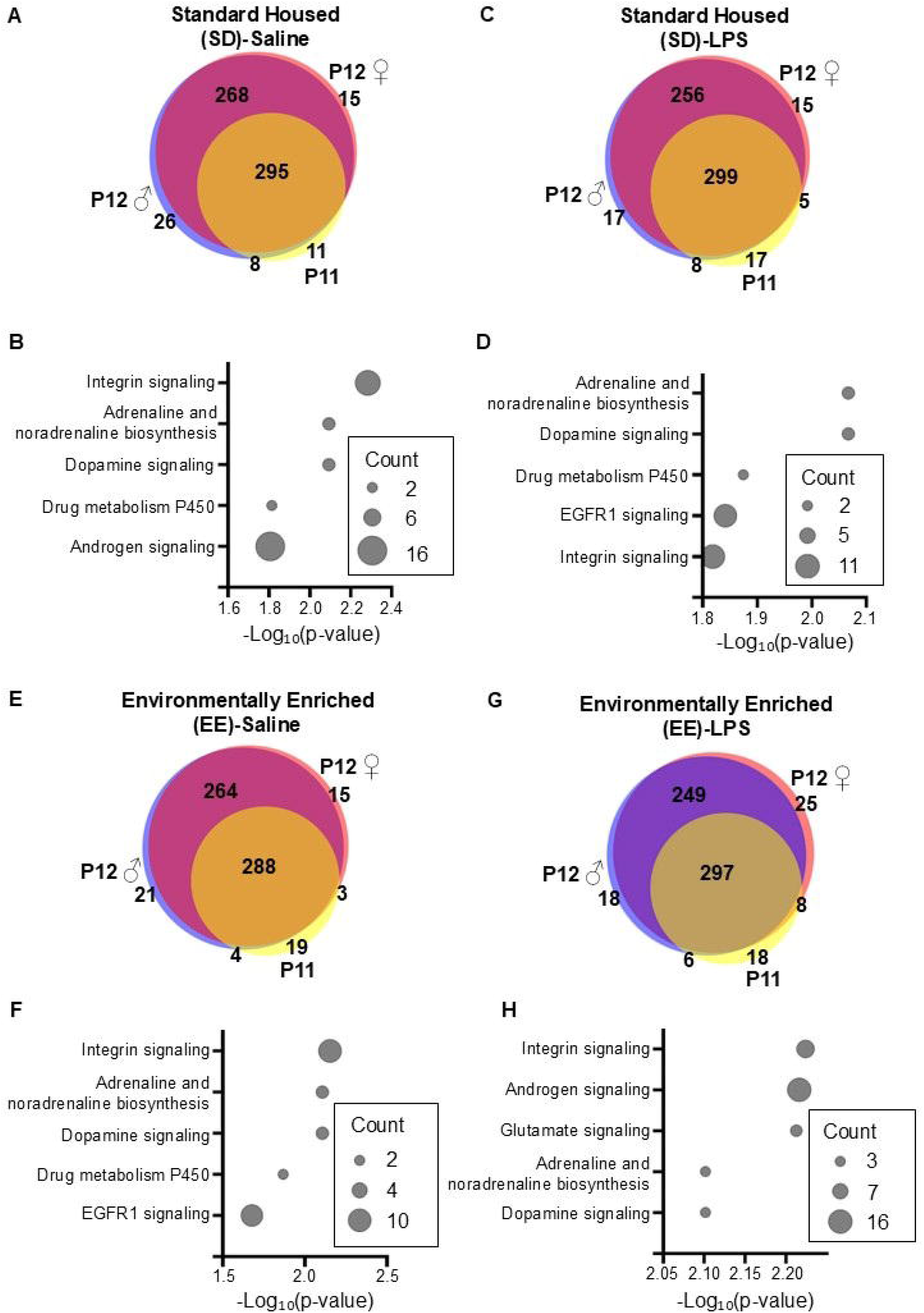
Overlapping miRNAs matched between maternal milk-derived extracellular vesicles (MEVs) and the hippocampus of their nursing male and female neonatal offspring. (**A, C, E, G**) Venn diagrams showing the number of miRNAs found in MEVs at P11 (yellow) and the hippocampus of male (blue) and female (red) offspring at P12, and their overlapping miRNA in groups (**A**) SD-Saline, (**C**) SD-LPS, (**E**) EE-Saline, and (**G**) EE-LPS. (**B, D, F, H**) The bubble plots show the -LogLL (p-value) of the top 5 enriched miRWalk pathways identified for the overlapping miRNA in (**B**) Figure A, (**D**) Figure C, (**F**) Figure E, and (**H**) Figure G, using the miRNA Enrichment Analysis and Annotation Tool (ccb-compute2.cs.uni-saarland.de/mieaa). miRNA count per pathway is indicated by bubble size. LPS: lipopolysaccharide; P: postnatal day; SD: standard housing; EE: environmental enrichment; n = 6.

## Discussion

Postpartum infections are known to affect breastfeeding people, however the consequences of illness on lactation and associated offspring outcomes are understudied^38^. Despite awareness of the importance of maternal health in the postpartum period, interventions aimed at supporting caregivers are often underutilized and undervalued. Compounding this gap is a limited understanding of the biological mechanisms through which such interventions may exert their effects. In this context, we identify MEV-miRNAs as a dynamic signaling pathway capable of conveying both stress-induced and protective environmental cues, from mothers to their offspring, during the lactational period. This highlights a potential mechanistic link between maternal experience and early developmental programing (see **Figure 6** for summary of proposed mechanisms).

**Figure 6.**
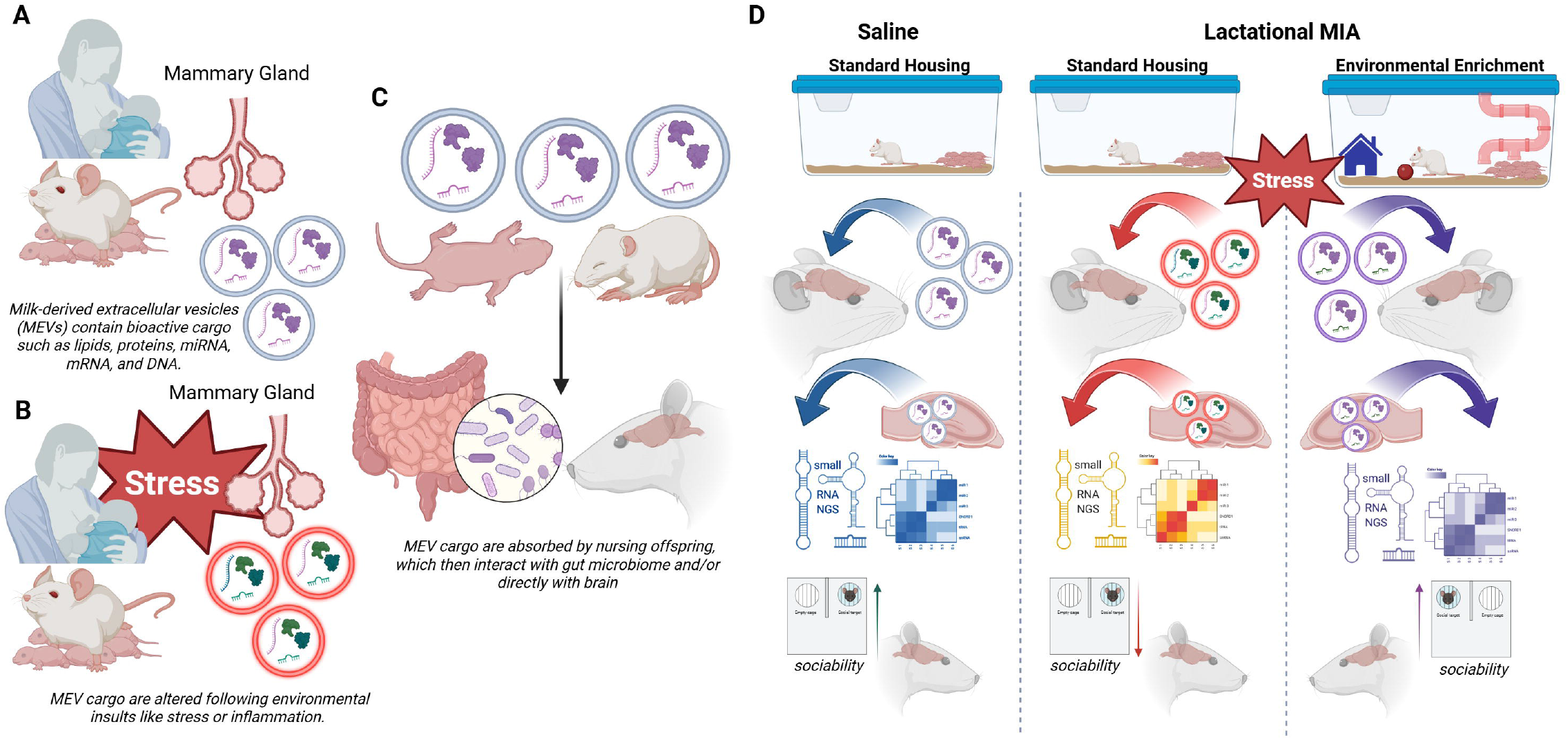
Overarching hypothesis of milk-derived extracellular vesicles (MEVs) as a mechanism of postnatal neuroimmune programming. (**A**) Nursing mammals pass MEVs through their milk to their feeding offspring. (**B**) Insults such as inflammatory stressors change the composition of MEV cargo. (**C**) Data suggest that when absorbed by nursing offspring, MEVs can interact with the gut microbiome and/or cross into and accumulate in the offspring brain. (**D**) We propose that maternal immune activation (MIA) alters the composition of MEV cargo, which then infiltrates the feeding offspring. Through either direct (brain) or indirect (e.g., gut microbiome) interactions, the altered MEV cargo (e.g., miRNAs), lead to transcriptional changes in brain regions like the hippocampus, affecting later life behavior in offspring. Environmental enrichment interventions can neutralize these changes, conserving behavior. Created in BioRender. Kentner, A. (2025) https://BioRender.com/50zpt07.

Consistent with prior work, we found that MIA increased corticosterone levels in maternal milk^39^, a glucocorticoid known to influence stress responsivity and emotional regulation in offspring^72.76^. However, EE housing did not attenuate MIA-induced elevations in milk corticosterone. This suggests that corticosterone is unlikely to mediate the protective effects of EE observed in this lactational MIA model. Moreover, while MIA caused a reduction in creamatocrit that was rescued by EE, offspring body weights were unaffected during the neonatal and weaning periods. Adult EE males did weigh more than their adult SD counterparts; however, the absence of the early weight differences, in combination with the early transcriptional and behavioral changes, supports the idea that other milk constituents may play a more prominent role in developmental programming.

Instead, our findings point to MEVs and their miRNA cargos as key mediators of environmentally driven signaling between mothers and nursing offspring. Here, MIA induced robust alterations in MEV miRNA cargo, particularly in mothers housed under standard laboratory conditions. In contrast, dams reared in EE displayed fewer MIA-induced changes in MEV-miRNA profiles, supporting the notion that enrichment stabilizes maternal biological outputs under stress. Notably, in MEVs from standard housed dams, novel miRNA-629 was consistently downregulated by MIA across both sampling days (P10 and P11), while novel miRNA-375 showed a biphasic expression pattern. These miRNAs predicted by miRDeep 2.0 and clustered by DAVID targeted genes involved in transcriptional regulation, ion channel function, and signal transduction, pathways critical to neurodevelopment and synaptic plasticity^77,78^. In MEVs from enriched dams, MIA upregulated novel miRNA-1347, which is predicted to target the expression of Tbl1xr1, a gene implicated in developmental disorders whose regulation confers protection^79-82^. This suggests that EE-modified MEV cargo programs developmental adaptation in the offspring via transcriptional processes. Importantly, the influence of the environment on MEVs is not transient.

The regulatory potential of MEV-miRNAs was consistent with changes observed in the neonatal hippocampus on P12. Indeed, LPS challenge resulted in a significant *downregulation* of miRNAs in the MEVs from SD mothers on P11, suggesting that there were fewer regulating mechanisms following MIA^28,29^. While it is also important to keep in mind the role of other MEV cargos (e.g., lipid and protein content), of the significant MEV-miRNAs in the EE housed dams there was more *upregulation*, suggesting that there may have been more regulatory mechanisms at play. Similar to the patterns of MIA-induced differentially expressed miRNAs in the MEVs, enrichment reduced the number of differentially expressed hippocampal miRNAs, indicating that EE can buffer against stress-induced transcriptomic modifications. Together, this suggests that EE housing stabilized the offspring brain during MIA through MEV delivery. Several miRNAs were expressed in both MEVs and the neonatal hippocampus, suggesting potential transfer of regulatory signals from milk to brain. Moreover, miR-709 and miR-128-1-5p were differentially expressed in neonatal hippocampus and showed the expected directional changes in their predicted gene targets (*Thbs1* and *Naa11*) in the adult hippocampal transcriptome. Cross-tissue miRNA-mRNA targets provide compelling support to the hypothesis that MEVs are sensitive to maternal environmental conditions and may relay this information to the developing offspring brain^7^.

Lactational MIA induced behavioral alterations in offspring that were also buffered by enrichment. In female offspring, MIA reduced the percent of time animals explored the center of the open field arena, indicative of heightened anxiety-like behavior; in male offspring, social preference was diminished by the MIA challenge. Importantly, EE exposure prevented these adverse outcomes in both sexes, consistent with prior studies demonstrating the protective effects of enriched housing on offspring stress regulation and social engagement^48-50, 83^. Interestingly, differences in maternal care were not observed across groups, suggesting that EE protection was not mediated by changes in overt maternal behaviors, such as time spent on the nest or frequency of licking/grooming. This is noteworthy, as enriched housing can modulate maternal care under non-stress conditions^51^, but the added “injection stress” from administering either LPS or saline during lactation may have masked group differences. Maternal care is sensitive to laboratory manipulations, and these findings caution against assuming that vehicle injections represent a neutral control. It is possible that relatively higher levels of parental support or enrichment may be needed, even in higher resource settings, to buffer against acute stressors. In general, it appears that the behavioral buffering effects of EE may instead involve more subtle or molecular-level changes in maternal signaling via MEVs in this lactational MIA model, rather than direct maternal care interactions.

In addition to sex-specific behavioral phenotypes, following lactational MIA, there were significantly different miRNA profiles in male and female hippocampus. The differential expression of MIA-induced hippocampal miRNAs in SD offspring was dramatically greater in males than in females, aligning with previous work showing sex-specific vulnerability to early life stressors^84^. While SD-LPS females had fewer changes in miRNA expression, it was notable that miR-1247-5p was differentially expressed in the hippocampus of this group. In vivo studies have identified this novel miRNA as neuroprotective following injury, reporting that it affected developmentally regulated biological processes such as differentiation, transcription regulation, and mitochondrial function^85^. This raises the possibility that females possess unique compensatory mechanisms that are activated under stress.

We acknowledge that there are limitations one should take into account when interpreting some of the findings. First, the dynamic nature of milk composition across the lactational period limits the temporal generalizability of our results. Our study only sampled across two days, and it remains unclear whether certain phases of lactation may be more or less responsive to either MIA or EE. Second, although we identified associations between miRNAs in MEVs and neonatal hippocampus, direct mechanistic evidence for MEV transfer and functional integration in the offspring brain is lacking. It is not known whether neonatal hippocampal miRNA profiles were directly reflective of deposited maternal MEV cargo or rather due to indirect actions between MEVs and the microbiome, influencing the hippocampal transcriptome, for example. Future studies using fluorescently labeled MEVs and individual miRNA cargo, in addition to MEV depletion and supplementation approaches are needed to directly test the uptake and influence of MEVs in brain. MEVs themselves carry multiple types of cargo, including lipids and peptides that can influence neurodevelopment, in addition to the miRNAs evaluated here. Finally, while our study highlights MEVs as plausible mediators of environmental signaling, we do not exclude interactive effects with other nutritive and non-nutritive milk constituents such as lipids, peptides, and glucocorticoids^39^, or factors such as pup separation which could have contributed to MEV-miRNA programming. The complexity of milk composition argues against a single-pathway explanation, and it is likely that additive or synergistic interactions across multiple milk and environmental factors are shaping neurodevelopmental trajectories.

## Conclusions

Breastfeeding remains the recommended source of nutrition during most maternal illnesses, as many pathogens including influenza, COVID-19, and foodborne agents are not transmitted directly through breast milk, yet antibodies against these pathogens are transmitted to offspring^86^. Critically, maternal illness and stress can alter milk composition, including the regulatory cargo of MEVs. Our findings demonstrate that MEV-miRNAs are sensitive to the maternal environment and can shape offspring neurodevelopment, highlighting the need for increased support for breastfeeding individuals. This includes a need for improved education and communication pathways to ensure breastfeeding individuals are informed and supported^87^. Indeed, enrichment exposure stabilized MEV cargo and mitigated the effects of MIA on offspring, positioning MEV-miRNAs as dynamic programming signals by which both stress-induced and protective cues are relayed. Given evidence linking breastfeeding to reduced neurodevelopmental disorder risk^88^, our study positions MEVs as a key mechanism by which maternal experiences influence infant brain development, with important implications for maternal-infant health policy and care.

## Supporting information

Supplemental Methods and Experimental Overview

## Funding and Disclosures

This project was funded by NIMH under Award Number R15MH114035 (to ACK), Touro Presidential Research Development Grant (to JM), the Manitoba Medical Service Foundation Grant (to SW), the Massachusetts College of Pharmacy and Health Sciences (MCPHS) Center for Undergraduate Research (TJL & ACK), a MCPHS Summer Undergraduate Research Fellowship (SURF) to TJL, an MCPHS New Discovery Award (ACK), and the Massachusetts Life Sciences Center – Capital Grants, which provided financial and equipment support. Electron Microscopy imaging and consultation services were performed in the HMS Electron Microscopy Facility (thank you to Maria Ericsson). We thank the Genomics Core Laboratory and its late Director, Dr. Brahmaraju Mopidevi, at the New York Medical College for the miRNA sequencing of the neonatal hippocampus and Mr. Marc Piquette for general laboratory support (MCPHS). The authors would also like to thank Azenta Life Sciences for their support with the miRNA sequencing of MEVs and adult hippocampal RNA sequencing. Figures 1, 2, 4, and 6 were made with BioRender.com and an earlier version of this manuscript was posted on the preprint server bioRvix. The content is solely the responsibility of the authors and does not necessarily represent the official views of any of the financial supporters. Sequencing datasets have been deposited to GEO (GSE300302, GSE300818).

## Author Contributions

Julia M, TJL, BH, DL, JS, JL, Jordan M, and ACK ran the experiments; Julia M., BH, Jordan M., BB, BB, SW, & ACK analyzed and interpreted the data; ACK wrote the manuscript; Julia M, BH, TJL, JS, JL, SW, Jordan M, & ACK edited the manuscript. Jordan M, SW, & ACK were involved in project design. Jordan M, SW, and ACK supervised trainees. ACK conceptualized the project.

## Conflict of Interest

The authors declare that they have no known competing financial interests or personal relationships that could have appeared to influence the work reported in this paper.

## Supplemental Methods and Experimental Overview

**Supplemental Table 1. Maternal immune activation reporting guidelines table**.

**Supplemental Figure 1. Maternal care and offspring outcomes following lactational maternal immune activation (MIA) and environmental enrichment (EE) housing**. Total distance traveled (meters) for (**A**) male and (**B**) female adult offspring. Maternal total time on nest (seconds) and (**D**) frequency of licking and grooming behaviors across P9 to P11. Adult body weights (g) for (**E**) male and (**F**) female offspring. Data are expressed as mean ± SEM; ##p <0.01, SD-LPS versus EE-LPS; LPS: lipopolysaccharide; P: postnatal day; SD: standard housing; n = 7-8.

**Supplemental Figure 2. Transcriptomic analyses of milk-derived extracellular vesicles (MEVs) 24-hours following maternal immune activation**. Volcano plots depicting the distribution of 2131 miRNA based on log2 fold change and –log10 p values in **A** SD-Saline compared to SD-LPS MEVs and **B** EE-Saline compared to EE-LPS MEVs. Each dot represents a miRNA, with blue representing downregulated miRNA and red representing upregulated miRNA. p < 0.05, FC > 1.3.

**Supplemental Table 2. Validation of miRNA-mRNA target predictions from nursing offspring and associated mRNA in the adult hippocampus**. The table shows the overlap of differentially expressed (p<0.01, FC>2) gene families (column) that are common in the hippocampus of male and female rats at P12 and present at P74 in their respective groups (rows). LPS: lipopolysaccharide; P: postnatal day; SD: standard housing; EE: environmental enrichment; n = 6.

**Supplemental Table 3. Statistical Reporting of Results**.

## Notes

### Competing Interest Statement

The authors have declared no competing interest.

### Summary of Updates

Figure, Figure 2, and Figure 4 have been revised to improve clarity. Revisions have been made to the methods, results, and discussion sections.

