## Supplemental Methods and Experimental Overview for "Investigating milk-derived extracellular vesicles as mediators of maternal stress and environmental intervention"

The animal protocols used in this study were approved by the Massachusetts College of Pharmacy and Health Sciences (MCPHS) Institutional Care and Use Committee and were carried out in compliance with the Association for Assessment and Accreditation of Laboratory Animal Care. **Figure 1A** illustrates the experimental timeline. Please see **Supplementary MIA Reporting Guidelines Table 1** (Kentner et al., 2019) and DeRosa et al., 2022a and 2022b for additional methodological details. Briefly, Sprague Dawley rats (Charles River, Wilmington) were housed and bred in either environmental enrichment (EE) or standard laboratory (SD) housing (DeRosa et al., 2022b; Núñez Estevez et al., 2020). As described in our prior work (DeRosa et al., 2022a), P10 rat dams (when milk is mature) were removed from their litter, weighed, and immediately administered either 100 μg/kg of the inflammatory endotoxin, lipopolysaccharide (LPS; Escherichia coli, serotype 026:B6; L-3755, Millipore Sigma), or pyrogen-free saline (intraperitoneal injection between 08:30hrs and 10:30hrs). Each dam was then placed into a clean standard laboratory “vacation” cage located in a separate procedure room to allow milk to accumulate for 2 hours. When separated from their dam, each litter remained together in a smaller clean cage that was positioned on a heating pad, to maintain body temperature, located in their regular housing room.

*Maternal behavioral evaluations*: MIA induction by LPS was validated by scoring the severity of sickness behaviors displayed by dams in their home cage. An observer blinded to the drug treatments scored measures of ptosis (droopy eyelids), piloerection (ruffled, unkempt coat appearance), and lethargy on a three-point scale (0 = none, 1 = mild, 2 = severe) as previously described (Connors et al., 2014) at baseline, 30, 120-, and 300-minutes post injection. A composite score was assigned for each time point (adapted from Hayley et al., 2002; Kentner et al., 2007). Maternal care was assessed in the morning (between 7:30am and 8:30am) and afternoon (between 3:00pm and 5:30pm; 60 minutes after neonatal huddling) on P9, P10, and P11. Dams were evaluated for 1-minute intervals over 6 observation periods on behaviors including the frequency of pup licking and grooming and the total time spent on the nest (DeRosa et al., 2022a; 2022b). The study group conditions for the dams were as follows: SD-Saline: n = 8; SD-LPS: n = 8; EE-Saline: n = 7; EE-LPS: n = 8.

*Milk collection methods and protein assays:* Two hours following their P10 saline or LPS injection, dams were anesthetized with isoflurane in O2 and administered 0.2 mL of oxytocin (20 USP/mL i.p.; 1OXY015/069241, Covetrus) to facilitate milk production (see DeRosa et al., 2022a; 2022b). Based on our milking protocol and their pharmacokinetic profiles, breastfeeding can resume immediately after anesthesia and/or the induced milk-let down response as neither isoflurane nor oxytocin are absorbed by the nursing offspring (Drugs and Lactation Database, 2020; Lee & Rubin, 1993; Par Pharmaceutical Inc, 2020). Moreover, our previous work demonstrated that although LPS-binding protein is detectable in maternal plasma, it does not transfer into milk following LPS challenge, indicating that the immunogen does not directly affect nursing offspring (DeRosa et al., 2022a). After cleaning the teats with distilled water, milk was collected by gently squeezing the base of each teat until the required milk volume was obtained from each dam (50-60 minutes for ~ 2.0 mL sample volume). Once dams fully awoke from the procedure (~5-10 minutes) they were returned to their litters in their regular home cage. On P11, dams were again separated from their litter (for 2 hours) followed by milk collection 24 hours after LPS or saline treatment (see DeRosa et al 2022a). Full litter body weights were evaluated prior to the dams return on P10 and P11. 20 µL of expelled milk from each dam was immediately collected into a microhematocrit tube, sealed, and placed into a hematocrit spinner (2 minutes at 13,700 x g; StatSpin^TM^ CritSpin Microhematocrit Centrifuge, Beckman Coulter, Inc). Following the procedures outlined by Paul et al., (2015) we calculated percent (%) creamatocrit based on sample separation into a cream versus clear layer (n = 7-8 dams/group). An additional 100 µL of each P10 sample was stored at -80°C for future evaluation of corticosterone (n = 7 dams/group). These samples were thawed and homogenized by overnight rotation on a Mini Tube Rotator (88861051, Fisher Scientific) at 4°C. The small sample assay protocol of the corticosterone ELISA kit (#ADI-900–097, Enzo Life Sciences) was followed, as recommended by the manufacturer, using a 1:40 dilution (DeRosa et al., 2022a; 2022b). In cases where values were below the assay's detection limit (e.g., in non-LPS-treated animals where corticosterone levels are typically low), standard pharmacological methods were employed, such as imputing undetectable values as zero, to facilitate statistical analysis (Barnett et al., 2021).

*MEV isolation:* The remaining milk sample from each dam was centrifuged at 3,000 x g for 10 min at room temperature (RT), to separate out the cream layer. The supernatant was collected and placed into a fresh tube and centrifuged again at 3,000 x g for 10 min at RT. The cream layer was removed immediately post collection and prior to ultracold storage of the clear supernatant. The differential ultracentrifugation combined with serial filtration (DUC) method (outlined in **Figure 2A**) is listed in the Minimal Information for Studies of Extracellular Vesicles (Welsh et al, 2023) guidelines as an acceptable approach for extracellular vesicle isolation. Indeed, DUC yields high quality MEV concentrations and is appropriate for biological experiments (Wijenayake et al., 2021; Wijenayake et al., 2024; Morgan et al., 2020; Chan et al., 2020). Following this approach, samples of clear supernatant, lacking cream, were thawed on ice and centrifuged twice at 3,000 x g for 10 min at 4°C. The supernatants were then centrifuged at 1,200 x g for 10 min at 4 °C (2x), at 21,500 x g for 30 min at 4 °C (2x), and once at 21,500 x g for 1 h at 4 °C, followed by filtration through 0.45 µm PES syringe filters (229749, Ultident) to clear fat globules and cellular debris. Casein proteins were precipitated using 17.5 molar acetic acid (1:100 (v/v)) and centrifuged at 4,500 x g for 30 min at 4°C as per Morozumi et al., 2021. The supernatant was filtered through 0.22 μm PES syringe filters (229747, Ultident) to remove carryover precipitates and remaining cell debris. To isolate MEVs, supernatants (now called the whey protein fraction) were ultracentrifuged at 100,000 x g for 120 minutes at 4°C using a Beckman Coulter XL-100 ultracentrifuge with a swing bucket SW55 TI rotor. The pellets containing enriched MEVs were washed in filtered 1X phosphate buffer solution (PBS) using a 1:1 (v/v of filtered PBS to whey protein fraction) and a second ultracentrifugation was completed at the same settings. The MEV pellet was then resuspended in filtered 1X PBS (1:2 v/v of filtered PBS to whey protein volume).

*MEV characterization:* All characterization and quantification of MEVs were in line with the MISEV 2023 guidelines using nanoparticle tracking analysis (NTA), western immunoblotting, and transmission electron microscopy (TEM; Welsh et al., 2023).

NTA: The concentration and particle size of isolated MEVs was determined with a NanoSight NS300 (Malvern Instruments, Ltd) using procedures adapted from Wijenayake et al., 2021 and Wijenayake et al., 2024. Samples were diluted with filtered 1X PBS. Data capture settings were adjusted to a detection threshold of 15, a camera level of 14, and three replicates of 25 second frame captures using a Green 532nm laser (**Figure 2B**).

Western blotting: Total soluble protein was extracted from MEVs by incubating samples on ice for 30 minutes in cell extraction buffer (FNN0011, Invitrogen) supplemented with protease inhibitor cocktail (1:20, v/v; PIC001.1, BioShop) and 1 mM phenylmethanesulfonyl fluoride (PMS123.5, BioShop), with intermittent vortexing. Lysates were centrifuged (13,000 rpm, 10 min, 4 °C), and protein concentrations were determined using the Pierce™ BCA Protein Assay Kit (23227, Thermo Scientific), per manufacturer’s instructions. Absorbance at 562 nm was measured with a BioTek Synergy H1 plate reader (BTSH1M2SI , Agilent Technologies). Samples were mixed with SDS loading dye containing 10% β-mercaptoethanol (SDS001.1; MERC002.500, BioShop), vortexed, denatured at 95 °C for 10 minutes, and stored at –20 °C.

Western immunoblotting was used to validate positive MEV markers such as Alix (E6P9B, Cell Signaling, 1:1000), Flotillin-1 (D2V7J, Cell Signaling, 1:1000), and CD9 (EXOAB-CD9-1, Systems Biosciences, 1:1000), in addition to the negative marker Calnexin (12186, Signalway Antibody, 1:1000). Protein lysates from MEVs were resolved on 8-12% SDS-polyacrylamide gels using a mini-Protean III electrophoresis system (165-3301, BioRad) with 15 µg of protein used for Alix and Flotillin-1, 20 µg used for CD9, and 50 µg of protein used for Calnexin. The gels were then transferred to a 0.45 µm PVDF membrane (IPVH00010, Millipore). Following a 1% casein block (v/v, 1X TBST) for 30 minutes and a 3×5 min wash in 1X TBST (or blocking in 5% casein solution in TBST for 1 hr at RT followed by 3 x 10 min washes in 1 X TBST for Calnexin), positive marker primary antibodies were incubated for 24 hours at 4°C on a shaker. Membranes were then washed as described above for each antibody and incubated with goat HRP-conjugated anti-rabbit IgG secondary (7074, Cell Signaling, 1 X TBST, 1:10,000 for 45 min for the positive markers and 1:5000 for 1 hour for Calnexin) at RT and again washed as described above for each primary. Visualization of membranes was done using the enhanced chemiluminescence ChemiDoc Imaging System (BioRad). The positive and negative MEV protein markers were not quantified to evaluate protein levels, rather it was determined whether the markers were either present or absent in the MEV samples and cellular controls (Wijenayake et al., 2021; Wijenayake et al., 2024; Welsh et al., 2023), authenticating MEVs in our isolated samples **(Figure 2C)**.

TEM: Adapting methods from Wijenayake et al., 2021, isolated MEVs were diluted with 1X PBS (1:10 v/v). 5µl of the diluted MEV sample was adsorbed for 1 minute to a carbon coated grid (CF400-CU, Electron Microscopy Sciences) that had been made hydrophilic by a 20 second exposure to a glow discharge (25mA). Excess liquid was removed with a filter paper (Whatman #1), the grid was then floated briefly on a drop of double distilled water (ddH_2_O to wash away phosphate or salt), blotted again on a filter paper (3x) and then negatively stained with 1% uranyl acetate (22400, Electron Microscopy Sciences) for 20-30 seconds at RT. After removing the excess reagent with filter paper, the grids were examined in a Tecnai™ G² Spirit BioTWIN transmission electron microscope and images recorded with an AMT Nanosprint 43-MKII camera at the Harvard Electron Microscopy Core Facility (**Figure 2D**).

*Neonatal offspring measures*: Offspring were evaluated on neonatal huddling (n = 7-8 litters/group), a form of social thermoregulation, 2 hours following the milking procedures on each of P10 and P11. Adapted from Naskar et al (2019), pups from each litter were equidistantly placed along the perimeter of a 40 cm x 40 cm arena and videotaped for 10 minutes. One video frame was extracted every 30 seconds, and the number of clusters (2 or more pups in physical contact) was averaged across 20 frames by blinded investigators (DeRosa et al., 2022a). On P12, 24 hrs following the last neonatal huddling recording, litters were weighed and one male and one female pup from each litter was euthanized with isoflurane, whole P12 hippocampus dissected, frozen on dry ice, and stored at −80°C. A single male and female from the remaining pups of each litter were weaned into same-sex pairs on P21, maintaining their original housing assignments.

*Adult offspring measures*: On P70, behavioral open field and social preference tests were evaluated following established methods (DeRosa et al., 2022a; 2022b; Núñez Estevez et al., 2020; n = 7-8 litters/group/sex). In the open field test, distance travelled (m) and the duration of time (seconds) spent in the center versus perimeter of an arena were recorded using automated tracking software (Any-maze, Wood Dale, IL). Avoidance behavior was quantified as the percentage of time spent in the center of the open field, calculated as [(time in center) / (time in center + time in perimeter)] × 100. After the habituation period in the open field, two cleaned wire containment cups were placed at opposite ends of the arena for a five-minute social preference test. One cup contained a novel, untreated standard housed rat of the same sex, age, size, and strain, while the other contained a novel object. The positions of the rat and the object were alternated across trials. Total time spent with each of the novel rat and object were evaluated by Any-Maze by blinded investigators. The social preference index was calculated for each subject using the formula: ([time spent with the novel rat] / [time spent with the novel rat + time spent with the object]) − 0.5 (Scarborough et al., 2020; DeRosa et al., 2022a; 2022b). On P74, animals were anesthetized with isoflurane and ventral hippocampus dissected, frozen on dry ice, and stored at −80°C. The procedures for whole hippocampus dissection and subsequent isolation of the ventral hippocampus are described by Benard and colleagues (2007). Briefly, after full hippocampal extraction, the two thirds in proximity to the retrosplenial cortex were identified as posterior/dorsal and the ventral hippocampus was identified as the remaining one third closest to the amygdaloid complex (Bernard et al., 2007; Nguyen et al., 2015; Strange et al., 2014) Our rationale for focusing on the adult ventral hippocampus stems from its role in mediating social interactions (Bagot et al., 2015; Felix-Ortiz and Tye, 2014) and anxiety (Adhikari et al., 2010, 2011), related to our behaviors of interest in this lactational MIA model.

*RNA extraction:* Total RNA was extracted from P12 and P74 hippocampal tissues using the RNeasy Lipid Tissue Mini Kit (74804, Qiagen) and from MEV samples using the Exosomal RNA Isolation Kit (58000, Norogen Biotek ), according to the manufacturer’s instructions. RNA samples were quantified using Qubit 2.0 Fluorometer (ThermoFisher Scientific) and RNA integrity was checked with 4200 TapeStation (Agilent Technologies).

*Small RNA-sequencing in MEVs (n = 6 dams/group)*: Small RNA sequencing library was prepared by using NEB Small RNA library Prep Kit (New England Biolabs). In brief, Illumina 3’ and 5’ adapter was added to RNA molecules with a 5’-phosphate and a 3’-hydroxyl group sequentially. A reverse transcription reaction was used to create single stranded cDNA. The cDNA was amplified via PCR using a common primer and a primer containing index sequence. Amplified cDNA construct was purified by polyacrylamide gel electrophoresis, and the correct band (~145 – 160 bp) excised from the gel and eluted with water. The eluted cDNA was concentrated by ethanol precipitation, to create the final sequencing library. The sequencing library was validated on the Agilent TapeStation (Agilent Technologies) and quantified by using Qubit 2.0 Fluorometer (ThermoFisher Scientific) as well as by quantitative PCR (KAPA Biosystems). The sequencing libraries were clustered on the flowcell. After clustering, the flowcell was loaded on the Illumina NovaSeq instrument according to manufacturer’s instructions. The samples were sequenced using a 2x150bp Paired End (PE) configuration. Image analysis and base calling were conducted by the Control software. Raw sequence data generated by the sequencer were converted into fastq files and de-multiplexed using Illumina's bcl2fastq 2.20 software. One mismatch was allowed for index sequence identification.

*Small RNA-sequencing in neonatal hippocampus (n = 6 litters/group/sex)*: Small RNA sequencing libraries were prepared by using Illumina TruSeq Small RNA Library Prep Kit (Illumina). Briefly, Illumina 3’ and 5’ adapters were added to RNA molecules with a 5’-phosphate and a 3’-hydroxyl group sequentially. A reverse transcription reaction was used to create single-stranded cDNA. The cDNA was then amplified via PCR using a common primer and a primer containing the index sequence. Amplified cDNA construct was purified by using Bluepippin, and the correct band (~145 – 160 bp) was eluted in a 40 µl fresh elution buffer. The sequencing library was validated on the Agilent TapeStation 4200 (Agilent Technologies) and quantified by using Qubit 2.0 Fluorometer (Invitrogen). The sequencing libraries were then multiplexed and sequenced on Illumina NextSeq 2000 in a 1x50bp configuration.

*RNA-sequencing in adult ventral hippocampus (n = 6 litters/group/sex)*: Strand-specific RNA sequencing library was prepared by using NEBNext Ultra II Directional RNA Library Prep Kit for Illumina following manufacturer’s instructions (NEB, Ipswich, MA, USA). Briefly, the enriched RNAs were fragmented for 8 minutes at 94 °C. First strand and second strand cDNA were subsequently synthesized. The second strand of cDNA was marked by incorporating dUTP during the synthesis. The cDNA fragments were adenylated at 3’ends, and indexed adapter was ligated to cDNA fragments. Limited cycle PCR was used for library enrichment. The incorporated dUTP in second strand cDNA quenched the amplification of second strand, which helped to preserve the strand specificity. The sequencing library was validated on the Agilent TapeStation (Agilent Technologies) and quantified by using Qubit 2.0 Fluorometer (ThermoFisher Scientific) as well as by quantitative PCR (KAPA Biosystems). The sequencing libraries were multiplexed and clustered onto a flowcell on the Illumina NovaSeq instrument according to manufacturer’s instructions. The samples were sequenced using a 2x150bp Paired End (PE) configuration. Image analysis and base calling were conducted by NovaSeq Control Software (NCS). Raw sequence data generated from Illumina NovaSeq were converted into fastq files and de-multiplexed using Illumina bcl2fastq 2.20 software. One mismatch was allowed for index sequence identification.

*Statistical procedures*: Three-way repeated measure ANOVAs (Housing x MIA x Time) were used to evaluate milk and behavioral endpoints, as appropriate. Two-way ANOVAs (Housing x MIA) were used for all other measures unless there were violations to the assumption of normality (Shapiro-Wilk test) in which case Kruskal-Wallis tests were employed (expressed as *X^2^*). To assess differences in the proportion of samples with detectable P10 milk corticosterone concentrations, the Fisher-Freeman-Halton exact test was used to appropriately account for undetectable levels in the SD-Saline group. The partial eta-squared (*n_p_*^2^) is reported as an index of effect size for the ANOVAs (Miles & Shevlin, 2001). To address sex as a biological variable (Clayton et al., 2018), both male and female animals were included and separate analyses were run for each sex (Ordoñes Sanchez et al., 2021; DeRosa et al., 2022a; 2022b). All data are expressed as mean ± SEM.

*Bioinformatic analysis*: (MEV miRNA-seq): Raw sequence reads were read trimmed to remove low-quality bases and adapter sequences using Trimmomatic (v0.30). Trimmed reads with a length of 18 to 32 bp were retained. Trimmed reads were compared to, and annotated with, the small RNA database (miRbase 22, [www.mirbase.org](http://www.mirbase.org)). For novel microRNA prediction, sequences were aligned to the corresponding genome and subjected to RNA folding and secondary structure analysis using miRDeep2 (v2_0_0_7). For quantified miRNAs based on hit counts, Miranda (v3.3a) was used to predict target sites based on microRNA sequences and the corresponding genomic cDNA sequences. Differential expression analysis was performed using the Bioconductor package edgeR (v3.4.6). Small RNAs with a p<0.01 and fold-change of normalized expression values >2 were considered significant for differential expression. (Neonatal hippocampus miRNA-seq): The raw sequence reads were quality, adapter trimmed and quantified using quantify miRNA tool in CLC Genomics workbench 20.0.4. Trimmed reads with a length of 15 to 31 bp were retained. Trimmed reads were compared to, and annotated with, the small RNA database miRbase 22. Hit counts for each small RNA were used as expression values and quantile normalization and log2-transformation were performed. Differentially expressed annotated small RNAs were identified via empirical analysis using the Fisher’s exact test. Differentially expressed miRNAs were considered statistically significant if p<0.01 and fold-change of normalized expression values >2. Benjamini–Hochberg false discovery rate corrected (FDR) p<0.05 and absolute fold-changes of normalized expression values >2 was used to find differentially expressed miRNAs that met this level of statistical significance. The miRNA-mRNA targets (target score>80) were predicted using miRDB (<https://mirdb.org/>) or miRDeep2 for Novel miRNAs. (Adult ventral hippocampus RNA-seq): Raw sequence reads were read trimmed to remove low-quality bases and adapter sequences using Trimmomatic (v.0.36). The trimmed reads were aligned to the genome of *Rattus norvegicus* (Rnor6.0 reference genome available on ENSEMBL) using the STAR aligner v.2.5.2b. Unique gene hit counts were calculated by using feature Counts from the Subread package v.1.5.2. Only unique reads that fell within exon regions were counted. DESeq2 workflow combined with Wald test was used for differential gene expression analysis to generate p values and log2 fold changes. Genes with p<0.01 and absolute fold-changes of normalized expression values >2 were considered differentially expressed for each comparison. Principal Component Analysis was performed using the plotPCA function of the DESeq2 R package to visualize the data, allowing for the identification of outliers, clusters, and relationships between samples. Heatmaps were generated using Multiple Experiment Viewer (MeV). Venn diagrams were generated using DeepVenn (<https://www.deepvenn.com>). Gene ontology was determined using the Database for Annotation, Visualization and Integrated Discovery (DAVID) functional annotation cluster tool (<https://david.ncifcrf.gov/>), while miRNA Enrichment Analysis and Annotation Tool (miEAA) was used to functionally annotate the different sets of miRNAs. Bubble plots and pie charts were generated in GraphPad Prism 10.4.2.
