## Supplemental Table 1 for "Investigating milk-derived extracellular vesicles as mediators of maternal stress and environmental intervention"

### Maternal Immune Activation Model Reporting Guidelines Checklist

| ARRIVE Reporting Guideline & Recommendation | Arrive Item | MIA Model Specific Reporting Recommendation<br>*Please complete this chart for each point outlined below. If not applicable, write N/A* |
|---|---|---|
| **Study design**<br>➢Overview of immune activation issues<br><br>For each experiment, give brief details of the study design including:<br>a. The number of experimental and control groups.<br>b. Any steps taken to minimize the effects of subjective bias when allocating animals to treatment (e.g. randomization procedure) and when assessing results (e.g. if done, describe who was blinded and when).<br>c. The experimental unit (e.g. a single animal, group or cage of animals).<br><br>A time-line diagram or flow chart can be useful to illustrate how complex study designs were carried out. | 6 | MIA Specific Reporting:<br>a. General need for improved reporting in MIA model methods + reporting pilot data<br>○ Details on pilot data:<br>a. Please see Figure 1 for experimental timeline outlining conditions.<br>b. LPS and Saline treated dams were matched to the same male for breeding.<br>c. The experimental unit is per litter - only one male and one female from each litter was used on any endpoint.<br>d. Offspring from saline and LPS dams were housed together at weaning. |
| **Experimental procedures**<br>➢Compounds<br>➢Validation measures<br><br>For each experiment and each experimental group, including controls, provide precise details of all procedures carried out. For example:<br>a. How (e.g. drug formulation and dose, site and route of administration, anaesthesia and analgesia used [including monitoring], surgical procedure, method of euthanasia). Provide details of any specialist equipment used, including supplier(s).<br>b. When (e.g. time of day).<br>c. Where (e.g. home cage, laboratory, water maze).<br>d. Why (e.g. rationale for choice of specific anaesthetic, route of administration, drug dose used). | 7 | Provide details of:<br>a. Compounds – source, vehicle, preparation/storage, administration route, volume administered, whether anesthetics were used at time of immune challenge.<br>○ Name of compound: lipopolysaccharide (LPS); Escherichia coli, serotype 02[illegible]<br>○ Catalogue number: L8274<br>○ Lot number: 093M4069V<br>○ Vehicle control used: pyrogen-free saline<br>○ Route of administration: i.p.<br>○ Volume administered: 100 ug/kg<br>○ Storage conditions: 4 degree Celsius fridge<br>○ Anesthetic (type, dose, duration) used:<br>Euthanized by isoflurane overdose<br>b. Housing variables at injection - temperature of room at injection time, cage change at time of injection or not<br>○ Light cycle of animal housing room: 0700-1900 Light<br>○ Time of day of injection: morning<br>○ Room temperature at injection time: RT<br>○ Did a cage change occur at time of injection: YES |

Kentner AC, Bilbo AD, Brown AS, Hsiao EY, McAllister AK, Meyer U, Pearce BD, Pletnikov MV, Yolken RH, Bauman MD. (2018). Maternal immune activation: reporting guidelines to improve the rigor, reproducibility, and transparency of the model. Neuropsychopharmacology, https://doi.org/10.1038/s41386-018-0185-7.

| | | |
|---|---|---|
| | | c. Validation of immune activation – behavior, physiological indices and/or cytokine data, including pilot dosing data<br>○ Method used to verify immune activation:<br>LPS treated dams had significantly higher passive home cage sickness behavior evaluations compared to saline dams.<br><br>d. Validation of gestational timing – vaginal plug, estrous cycle, weight gain<br>○ Method of validating gestational timing:<br>Constant body weight gain.<br><br>Additional comments: |
| **Experimental animals**<br>➢Species/strain/vendor<br><br>a. Provide details of the animals used, including species, strain, sex, developmental stage (e.g. mean or median age plus age range) and weight (e.g. mean or median weight plus weight range).<br>b. Provide further relevant information such as the source of animals, international strain nomenclature, genetic modification status (e.g. knock-out or transgenic), genotype, health/immune status, drug or test naïve, previous procedures, etc. | 8 | Provide details of:<br>a. Species – considerations for appropriate species (mouse, rat, non human primate, other)<br>○ Species: Rat<br><br>b. Strain – variability in strain can influence model<br>○ Strain: Sprague-Dawley<br><br>c. Maternal/Offspring Physiological Variables at time of immune challenge – age, body weight<br>○ Maternal Age at challenge: unknown - ordered according to body weight (bet<br>○ Maternal Body weight: ~350 grams<br>○ Offspring Age at challenge: offspring not challenged, but dams were challeng<br>○ Offspring Sex: Both males and females tested<br>○ Offspring Body weight: Full litter weight at postnatal day 10: ~270g.<br><br>d. Vendor – even within the same strain, vendor can influence endpoints<br>○ Vendor: Charles River<br>○ Location of Vendor: Willimington, MA<br>○ Room/area where animals originated from: Females: K98; Male |

| | | |
|---|---|---|
| | | Additional Comments:<br>N/A |
| **Housing and husbandry**<br>➢Cage, ventilation, bedding, enrichment<br><br>Provide details of:<br>a. Housing (type of facility e.g. specific pathogen free [SPF]; type of cage or housing; bedding material; number of cage companions; tank shape and material etc. for fish).<br>b. Husbandry conditions (e.g. breeding program, light/dark cycle, temperature, quality of water etc for fish, type of food, access to food and water, environmental enrichment).<br>c. Welfare-related assessments and interventions that were carried out prior to, during, or after the experiment. | 9 | Provide details of:<br>a. Caging systems<br>○ *At breeding*<br>Material of cage: standard housing: polycarbonate; Enriched housin<br>Cage dimensions: standard housing: 51 x 41 x 22 cm; Enriched hou<br>○ *After parturition*<br>Material of cage: standard housing: polycarbonate<br>Cage dimensions: just before parturition standard housed: 27 x 48 x 2<br>○ *At weaning*<br>Material of cage: standard housing: polycarbonate; Enriched housir<br>Cage dimensions: standard housed: 27 x 48 x 20 cm; Enriched sepa<br><br>b. Animal Holding room<br>○ Temperature in room: RT (~21.Celsius)<br>○ Humidity in room: ~40<br>○ Ventilation system: HVAC; open topped cages<br>○ Specific pathogen free [SPF]: NO<br>○ Are males & females housed in the same or separate rooms:<br>Housed in same room after weaning<br>c. Bedding exchanges/bedding type<br>○ Type of cage bedding used: bed-o'-cobs Laboratory Animal Bedding by The<br>○ Frequency of cage changes per week<br>*during gestation:* twice weekly<br>*during neonatal period:* one time weekly<br>*following weaning:* twice weekly<br><br>d. Breeding - bred on site or timed pregnant, how many different sires (are the same fathers breeding with both experimental and control dams)<br>Breeding location: on site |

- Gestational age at shipping: Shipped prior to breeding
- Biological age of dams (if not listed in Section 8c): unknown; shipped
- Number of Dams bred: 31
- How many times have dams been mated previously: 0
- How many times did the dams mate and not become pregnant: 1 time
- Are the dams primiparous or multiparous? All dams are primiparous
- What was the frequency of maternal handling during the gestational/neonatal period (e.g. cage cleanings, weighing, blood collection manipulations): once or twice weekly cage changes (see above),
- Biological age of sires: specific age unknown, go by weight (250-275 grams
- Number of sires bred: 10
- How many times have sires been mated previously: 1-4 times
- How many times did the sires mate successfully (e.g. mating resulted in pregnancy, full term birth): 1-4 time
- If bred previously, what was the interval between mating times:
- Are sires matched to experimental and control dams: YES
- Describe the mating design (1:1, 1:2 etc): 1 male to 2 females

e. Social enrichment – number of cage companions
- Number of cage companions prior to breeding: 2
- Gestational age when dam separated for parturition: gestational day 19
- Number of cage companions at weaning: 2 animals housed per cage

f. Physical enrichment – describe enrichment devices, and when enrichment is in the cage (removed when pups born? Or present throughout study), does the enrichment type change? How frequently?
- Describe what type of enrichment devices (and how many) are included in cage/housing room:

Standard caging system: access to one tube, one chew bone and Nestlets® (Ancare, Bellmore, NY). Bone and tube removed on gestational day 19 and replaced at postnatal day 14. Nestlets® available throughout study.

Enrichment caging system: access to a ramp, some tubes, 3-5 toys (e.g. plastic cars, standard pet toys, ropes, things to climb); toys changed out and replaced twice weekly to increase novelty

| | | |
|---|---|---|
| | | ○ Does enrichment type/access change across study? YES<br>○ If so, when does enrichment type/access change (e.g. enrichment removed prior to parturition and replaced in late neonatal period):<br>Tubes and chew bones removed on gestational day 19, for pup safety. Nestlets® available throughout study.<br>Enrichment litters receive enrichment back at postnatal day 14; toys continue to be replaced twice a week for novelty. Standard housed offspring given standard enrichment devices at weaning.<br>Additional Comments:<br>N/A |
| **Sample size**<br>➢Litter versus offspring<br>a. Specify the total number of animals used in each experiment, and the number of animals in each experimental group.<br>b. Explain how the number of animals was arrived at. Provide details of any sample size calculation used.<br>c. Indicate the number of independent replications of each experiment, if relevant. | 10 | Provide details of:<br>a. Maternal N vs offspring N<br>○ What is the total number of dams/litters included in the study: 32<br>○ What is the total number of offspring per litter included the study: 1 male and 1 female<br>b. Litter size and sex distribution<br>○ What size was each litter maintained at: 10 pups per litter<br>○ What age did culling take place at: postnatal day 1 (day of birth being<br>○ How many males and females were maintained in each litter: equal distribution, wherever possible<br>c. Cross fostering<br>○ Did cross fostering occur: NO<br>○ If so, at what age did cross fostering occur: N/A<br>Additional Comments:<br>N/A |

| | | |
|---|---|---|
| **Allocating animals to experimental groups**<br><br>a. Give full details of how animals were allocated to experimental groups, including randomization or matching if done.<br>b. Describe the order in which the animals in the different experimental groups were treated and assessed. | 11 | a. How many offspring per litter were used in each measure: entire litters were utilized<br><br>b. Randomization/Matching procedures<br>○ What procedures were used to assign animals to groups:<br>pseudorandomization<br><br>c. Sex as a biological variable (behavioral and physiological outcomes)<br>○ Were both males and females evaluated in each behavioral and physiological outcome: YES<br><br>Additional Comments:<br>N/A |
| **Experimental outcomes**<br>➢ Behavioral testing<br>➢ Physiological endpoints<br><br>Clearly define the primary and secondary experimental outcomes assessed (e.g. cell death, molecular markers, behavioral changes). | 12 | a. Maternal behavior and pup interactions<br>○ If maternal care was evaluated, were there differences following immunogen challenge (if so, please briefly describe):<br>See manuscript<br><br>b. Age(s) of offspring at behavioral testing/physiological evaluation endpoints:<br>see Figure 1 - for experimental outline and offspring age for each endpoint<br><br>c. Order of testing (e.g. behavioral test order)<br>○ Were animals evaluated in a counter-balanced order in terms of:<br>*presentation of tests to each animal:* NO<br>*order of experimental/control groups run through each test:* YES<br>○ What was the inter-test interval if a single animal underwent a battery of tests:<br>Open field and social preference occurred on same day. |

| | | |
|---|---|---|
| | | Additional Comments:<br>Presentation of tests were not counterbalanced because the open field test must occur prior to to the social preference test. |
| **Statistical methods**<br>a. Provide details of the statistical methods used for each analysis.<br>b. Specify the unit of analysis for each dataset (e.g. single animal, group of animals, single neuron).<br>c. Describe any methods used to assess whether the data met the assumptions of the statistical approach. | 13 | a. Unit of analysis for each data set<br>○ Is the unit (n) of each analysis based on number of litters, or number of animals used per group:<br>n = litter - - only one male and one female from each litter was used on any endpoint. |
| **Other Disclosures** | | Please make note of any other extraneous variables that you would like to report (e.g. fire alarms, construction, temporary relocations, other variables that you think we should be considering in our studies etc.):<br>N/A |

The recommended use of this reporting form is to fill it out and include it as supplemental material for each of your laboratory's research publications. If there are difficulties utilizing/adapting this fillable form, please contact one of the corresponding authors to request a copy. The authors give permission for this table to be edited for use in reporting on other animal models (e.g. postnatal immune challenge models, early life stress models) as appropriate.

Kentner AC, Bilbo AD, Brown AS, Hsiao EY, McAllister AK, Meyer U, Pearce BD, Pletnikov MV, Yolken RH, Bauman MD. (2018). Maternal immune activation: reporting guidelines to improve the rigor, reproducibility, and transparency of the model. Neuropsychopharmacology, https://doi.org/10.1038/s41386-018-0185-7.
