## Supplementary figures and images for "Investigating milk-derived extracellular vesicles as mediators of maternal stress and environmental intervention"

### Supplemental Figure 1

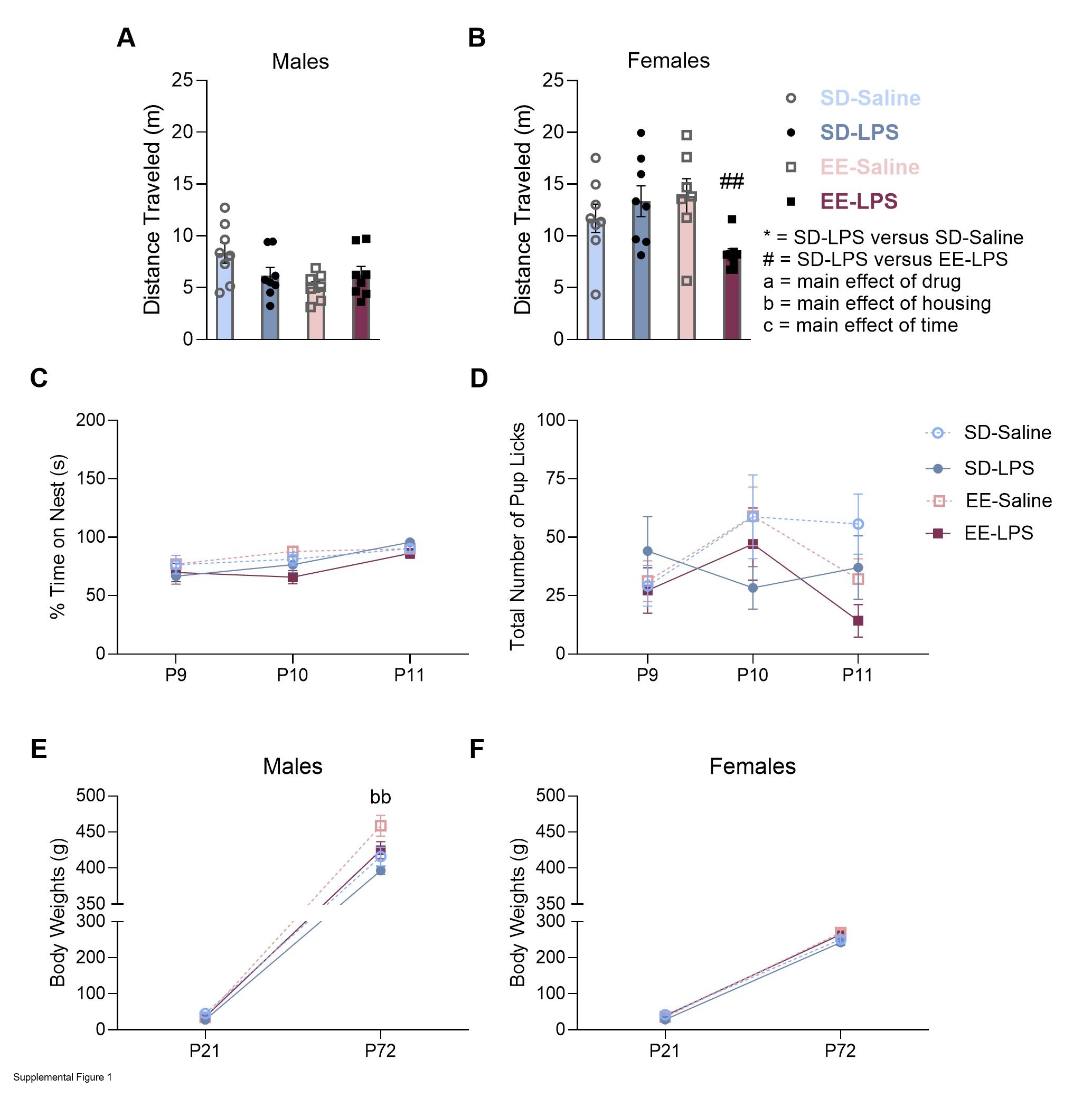

### Supplemental Figure 2

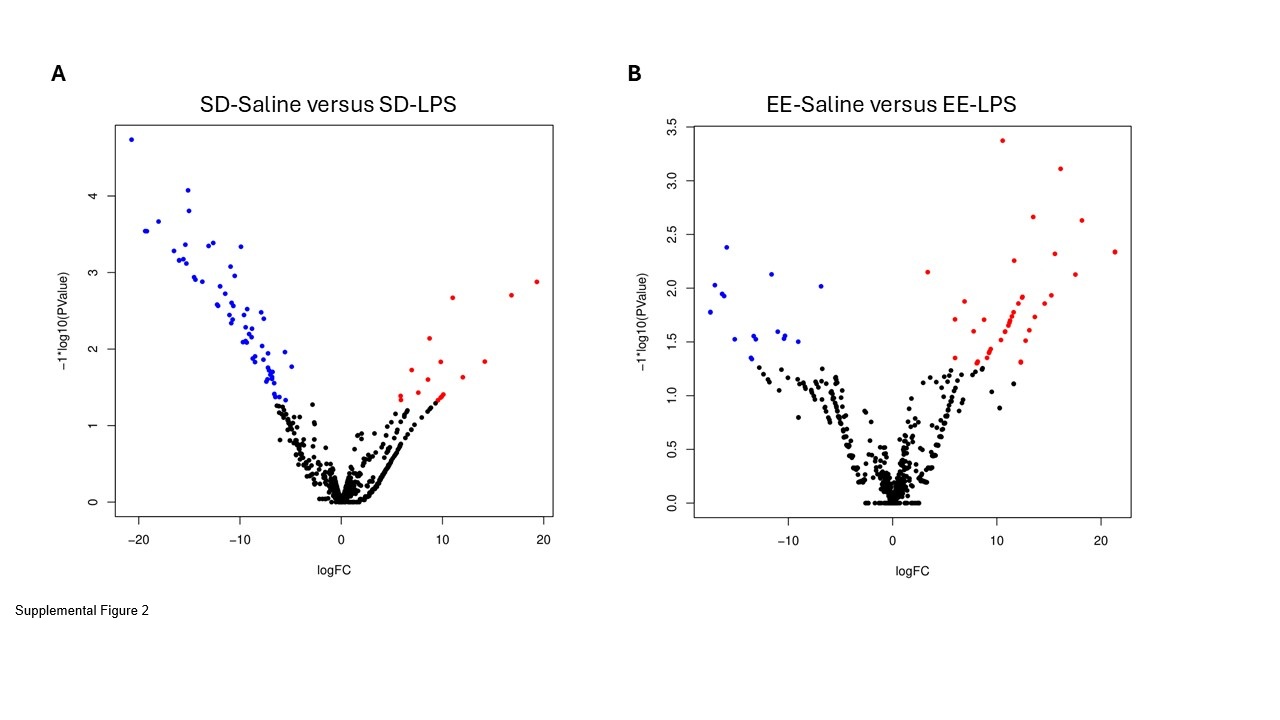
