## Supplemental Table 2 for "Investigating milk-derived extracellular vesicles as mediators of maternal stress and environmental intervention"

**Supplemental Table 2.** Validation of miRNA-mRNA target predictions from nursing offspring and associated mRNA in the adult hippocampus

| **Group** | **Gene family** |
| --- | --- |
| SD LPS vs. Saline ♂ | *Cd, Clec, Lrrc, Naa, Slc39a, Thbs, Vom2r* |
| SD LPS vs. Saline ♀ | *Sema, Slc, Ube2* |
| EE LPS vs. Saline ♂ | *Cdk/Cdkn* |
| EE LPS vs. Saline ♀ | *Col, Rab, Tmem* |
