## Supplemental Table 3 for "Investigating milk-derived extracellular vesicles as mediators of maternal stress and environmental intervention"

| **Endpoint** | **Statistical Test & Result** | **Figure** |
| --- | --- | --- |
| Maternal sickness behavior | *MIA x Housing x Time repeated measures ANOVA*:  EE and SD dams following lactational saline or LPS challenge; *MIA x Time interaction:* F(2,60) = 5.421, p = 0.007, ηp^2^ = 0.153. LPS treated dams had higher sickness scores, 5-hours post MIA challenge: *main effect of drug*: p<0.001.  *Housing x Time interaction*: F(2,60) = 4.574, p = 0.017, ηp^2^ = 0.175; SD dams appeared less healthy than EE mothers at baseline (*main effect of housing*: p<0.01), but EE did not protect against MIA-induced sickness (p>0.05). | Figure 1B |
| Maternal total time on nest (seconds) | *MIA x Housing x Time repeated measures ANOVA*:  MIA x Housing interaction: F(2, 56) = 3.890, p = 0.045, ηp^2^ = 0.122 for total time (seconds) that dams spent on the nest across P9 to P11, but no group differences were observed with post hoc analysis (p>0.05). | Supplemental Figure 1C |
| Maternal licking & grooming behaviors | *MIA x Housing x Time repeated measures ANOVA*:  There were no differences in the frequency of licking and grooming maternal behaviors across P9 to P11 (p>0.05). | Supplemental Figure 1D |
| Neonatal huddling behavior | *MIA x Housing x Time repeated measures ANOVA:*  EE protected against MIA associated reductions in offspring huddling behavior (average number of pup clusters) across P10 and P11 (*MIA x Housing interaction*: F(1, 29) = 5.003, p = 0.003, ηp^2^ = 0.147). SD-LPS versus SD-Saline: p <0.01; SD-LPS versus EE-LPS: p <0.05. | Figure 1C |
| Adult total distance traveled (m) | *MIA x Housing ANOVA:*  *MIA x Housing interaction* for male: F(1,27) = 4.266, p = 0.049 and female (F(1,27) = 7.522, p = 0.011) adult offspring. Female EE-LPS offspring traveled less than SD-LPS (p = 0.006) and EE-saline (p=0.005) females. | Supplemental Figure 1A, B |
| Adult open field test | *MIA x Housing ANOVA:*  Neither MIA nor housing affected adult male offspring in the open field test (*MIA x Drug interaction*: p>0.05), whereas SD-LPS female offspring spent less percent (%) time in the center of the open field, which was prevented by EE housing (*MIA x Drug interaction*: F(1,27) = 5.261, p = 0.030, ηp^2^ = 0.300); SD-LPS versus SD-Saline: p<0.01; SD-LPS versus EE-LPS: p<0.05. | Figure 1D,E |
| Adult social preference | *Kruskal-Wallis test:*  Male SD-LPS offspring had a significantly reduced social preference index (versus SD-Saline: *X^2^*(1) = 5.835, p = 0.016) that was buffered by EE (SD-LPS versus EE-LPS: *X^2^*(1) = 5.906, p = 0.015), while female offspring were not affected (p>0.05). | Figure 1F,G |
| Milk creamatocrit | *MIA x Housing ANOVA:*  Percent (%) creamatocrit was reduced in the milk of SD-LPS dams, which was mitigated by EE housing (*MIA x Housing interaction*: F(1, 29) = 16.923, p = 0.001, ηp^2^ = 0.369). SD-LPS versus SD-Saline: p <0.01; SD-LPS versus EE-LPS: p <0.05. Main effect of time: p <0.0001. | Figure 1H |
| Neonatal offspring full litter body weights (g) | *MIA x Housing x Time repeated measures ANOVA:*  The full litter body weights of nursing pups were not affected by either MIA or housing across P10-P12 (*MIA x Housing x Time interaction*: p>0.05). | Figure 1I |
| Milk corticosterone (ng/mL) on P10 | *Fisher-Freeman-Halton Exact test:*  Overall omnibus test*:* p = 0.001  SD-Saline versus SD-LPS: p = 0.001  SD-LPS versus EE-LPS: p > 0.05  EE-Saline versus EE-LPS: p = 0.15  SD-Saline versus EE-Saline: p > 0.015 | Figure 1J |
| Adult offspring body weights (g) | *MIA x Housing ANOVA:*  Housing x Time interaction: F(1, 24) = 8.051, p = 0.009, ηp^2^ = 0.546 for male offspring body weights (g). Follow up tests showed that EE males (440.46 ±10.073 g) were heavier than SD males (405.69±6.978 g) on P72 but no significant differences in female offspring body weights (p>0.05). | Supplemental Figure 1E,F |
| Total volume of MEVs recovered (uL) | *MIA x Housing x Time repeated measures ANOVA*  MIA x Time interaction: F(1, 20) = 14.414, p = 0.001, ηp^2^ = 0.419; a smaller volume of isolated MEVs was recovered from MIA milk on P11 which was not protected by EE; *main effect of MIA*: p<0.01. | Figure 2E |

*SD-Saline: n = 8; SD-LPS: n = 8; EE-Saline: n = 7; EE-LPS: n = 8. LPS: lipopolysaccharide; P: postnatal day; SD: Standard Housed; EE: Environmental Enrichment; MIA: Maternal immune activation.
